# Surrogate Gradients for Gradient-Based Parameter Estimation in Simplified Neuron Models

**DOI:** 10.64898/2026.09.19.752831

**Authors:** Paul Mayer, Alexander Kozlov

## Abstract

Simplified spiking neuron models such as the Adaptive Exponential Integrate-and-Fire (AdEx) model reproduce the firing behaviour of diverse neuron types at a fraction of the cost of biophysically detailed models, making them attractive for large-scale brain simulation. Fitting their parameters to recordings, however, still relies on derivative-free search such as grid search or evolutionary algorithms, because the discrete spike-and-reset mechanism renders these models non-differentiable. Surrogate gradients, used in deep spiking networks, replace the spike derivative with a smooth approximation on the backward pass. Whether they also enable efficient gradient-based parameter estimation for single simplified neurons is untested.

We implement a differentiable AdEx model with a surrogate-gradient variant in the Jaxley framework. The implementation is on par with target-compiled C++ Brian2 code while remaining end-to-end differentiable, vmap-batchable and GPU/TPU-portable. It is 200 times faster than Brian2’s default runtime mode.

Across a recovery-radius benchmark of 2,700 paired runs, gradient-based fitting never matched a gradient-free Nelder–Mead baseline. At sub-threshold levels, gradient descent instead beats Nelder–Mead. The obstacle is therefore the spike mechanism and the loss geometry it induces, not gradient descent itself.

## 1 Introduction

In the last century, studying and modelling neurons has been seen as the gateway of understanding the brain in its entirety. What first started with very simple Leaky Integrate-and-Fire (LIF)-models, quickly became proving ground for more complex and more sophisticated ideas. Lack of computational resources has been a primary driver for why simplified neuron models were still developed, because they promised to be the only way to really model parts of the brain on super-computers 20 years ago. With the steady increase in computing power, the research community shifted focus from those models in favour of more sophisticated Hodgkin-Huxley (HH)-type models, which require more computational resources to simulate and to fit parameters to. However, the idea of simulating the brain as whole has remained an unreachable goal for many decades, until now.

Computational power has reached whole new levels, and using computationally less intensive simplified models might be the key step to perform whole brain simulations. Adaptive models like Adaptive Exponential Integrate-and-Fire (AdEx) can faithfully capture the firing behaviour of diverse neuron types and therefore promise to be a prime candidate for realistic large-scale simulations. However, this needs very fast and efficient, reliable, and effective optimization methods to fit neuron models to experimental data. Due to the non-differentiable nature of simplified models, optimization methods like grid search or evolutionary algorithms are used instead of gradient methods.

In this work, we investigate whether surrogate gradient techniques can enable gradient-based parameter estimation for simplified neuron models. Specifically, we ask:

1. can surrogate gradients produce useful parameter updates for the AdEx model,
2. what loss function design is required to achieve successful optimization and
3. where does gradient-based fitting fail relative to derivative-free methods, and why?

To investigate this, we make the following contributions:

1. We implement a surrogate gradient-enabled AdEx model within the state-of-the-art Jaxley framework, enabling gradient-based parameter optimization.
2. We investigate loss function requirements and compare voltage-based Mean Squared Error (MSE), feature-based and distance-based losses.
3. We systematically test the robustness of the gradient-based optimization, measuring spike timing accuracy using a coincidence factor metric.
4. We show that gradient-fitting does not transfer to AdEx from differentiable HH-type models, and we address the potential failure modes.

Further work is needed to enable efficient and effective parameter optimization using gradient-based optimization.

In this work we bridge two worlds: gradient-based optimization and the simplified neuron models used in biophysical simulation. Section 2 gives the necessary background and section 3 surveys the related work. Section 4 describes how we optimize and evaluate the AdEx model: the loss functions we compare, the training setup, and the recovery-based benchmarks. It closes with our first contribution (section 4.3) — a differentiable AdEx model in `Jaxley`, together with the surrogate-gradient variant that makes gradient-based fitting possible. Section 5 reports our findings and section 6 interprets them. We conclude in section 7 with future work and ethical considerations.

## 2 Background

This chapter introduces the technical foundations required to understand gradient-based parameter fitting in simplified neuron models. We start with the Adaptive Exponential Integrate-and-Fire (AdEx) model and explain why its parameters need to be fitted (Section 2.1). We then discuss the landscape of parameter tuning methods, motivating why gradient-based approaches are attractive but face a fundamental obstacle: the non-differentiable reset mechanism (Section 2.2). Surrogate gradients overcome this obstacle, but introduce their own challenges — vanishing gradients through leak dynamics and sensitivity to the surrogate shape (Section 2.3). Finally, even with working gradients, the loss function determines whether optimization succeeds. We survey three categories of loss functions and their trade-offs for spiking models (Section 2.4).

### 2.1 Neuron Models

Large-scale brain models are limited in size mostly by computational feasibility. Detailed biophysical models which combine multi-compartment reconstructions of neuron morphology with non-linear ion channel dynamics (Hodgkin-Huxley (HH), stochastic models) can precisely predict membrane behaviour of a vast variety of neuron types. However, they require huge computational resources to estimate the required parameters to fit the models to experimental data. Simplified Neuron Models overcome this challenge, capturing a subset of the sophisticated neuron behaviour with only a fraction of the computational cost, enabling large-scale simulations [1]. Point neuron models boost computational performance by reducing branched neuron morphologies to a single active compartment. The AdEx model [2] provides a particularly interesting balance — it reproduces a variety of firing patterns but drastically reduces the number of necessary parameters of the model. This makes it a favourable model for large-scale neuron simulations [3].

The AdEx model improves the simpler Leaky Integrate-and-Fire (LIF) model in two ways: first, it introduces exponential behaviour when initializing the spike. Second, the model allows for adaptation, meaning it can change spiking frequencies or post-spike refractoriness based on their firing history. It is defined by the equations given in [2, 4]:

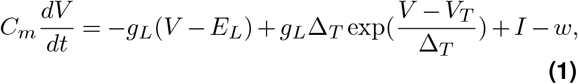

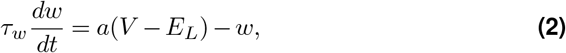

with the following reset condition:

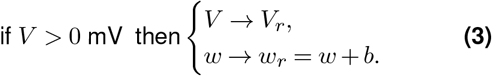

It describes the membrane potential over time *V* (*t*) given an injected current *I*(*t*). Equation (1) describes the change of potential. *C* is the membrane capacitance, *g*_*L*_ the leak conductance and *E*_*L*_ the effective resting potential. We can understand −*g*_*L*_(*V*− *E*_*L*_) as the leak current that slowly pulls the membrane potential towards its resting potential. 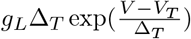 models the depolarization initialized by the fast reacting sodium channels using an exponential function, modelled using a driving force based on an effective threshold potential *V*_*T*_ and a slope factor Δ_*T*_. *I* is the injected current, and *w* is the adaptation current, which is described in equation 2.

The adaptation mechanism is controlled via an adaptation current which opposes depolarization. Unlike the fast membrane behaviour, this current is related to the time constant *τ*_*w*_, which controls the decay time. It is controlled by *a* (subthreshold adaptation) as well as *b* (spike-triggered adaptation).

Even though the number of parameters is significantly reduced compared to biophysical HH-type models, it still requires parameter fitting — especially because some parameters (e.g. *a, b* or *τ*_*w*_) do not have direct physiological counterparts (like *C* or *g*_*L*_). For AdEx to perform as an effective tool, efficient methods for parameter optimizations are essential.

The AdEx equations (Eqs. 1,2) are continuous ordinary differential equations with a discontinuity at each spike event. To simulate numerically, we discretize into a series of time steps *t*_0_, *t*_1_ = *t*_0_ + Δ*t*, …, *T*. Given the state *s*_*t*_ = (*V*_*t*_, *w*_*t*_) at time *t*, we compute *s*_*t*+1_ using forward Euler for *V* and exponential Euler for *w*. This produces a sequence of states *s*_0_, *s*_1_, …, *s*_*T*_ — the same that Backpropagation Through Time (BPTT) (Section 2.3.1) will differentiate through. It is important to note that both integration method and time step size Δ*t* affect both simulation accuracy and gradient quality.

### 2.2 Parameter Tuning for Neuron Models

A central challenge in neuroscience has been to ‘identify the parameters of detailed biophysical models, such that they match physiological measurements at scale’ [5] in reasonable simulation runtimes. The parameter space is highly non-convex, which is a challenge for any optimization approach. We further dive into this problem in Sec. 2.4.1. The zoo of optimization techniques is large, spanning from evolutionary algorithms [6] to gradient descent [7], however, deciding which algorithm to use may often depend on the underlying properties of the model. For tuning simplified neuron models, many different approaches have been used.

#### 2.2.1 Non-gradient-based Tuning

Derivative free methods like Nelder–Mead have successfully been used to fit parameters. It has simple design principles: it maintains a simplex (geometric shape of *n* + 1 vertices in *n* dimensions) and iteratively transforms it through four operations: reflection, expansion, contraction and shrinkage. At each step, the worst vertex is replaced by a better point found through these geometric operations. The simplex adapts its shape to the landscape — elongating along valleys, shrinking near minima, ending when it becomes sufficiently small. Parameter optimization is well-suited in regards to low-dimensionality: neuron models don’t have many parameters which is good (5-12); however, it’s a local optimizer. Simplex algorithms converge to the nearest minimum from its starting point. There exist no mechanism to escape local minima in non-convex landscapes, which requires lots of iterations with different starting positions.

A different popular choice are Evolutionary Algorithms (EAs): These models maintain a collection of candidate solutions (population, parameter sets). At each generation, fitness is evaluated using an objective functions. Fittest solution ‘produces offspring’ through mutation and recombination. Similar to the simplex algorithm, this only requires forward evaluation of the simulator. Multi-objective variants optimize for several objective functions simultaneously. This is a useful feature because for neuron fitting, multiple features (spike count, timing, frequencies, subthreshold voltage, …) extracted from multiple traces can be treated as separate objectives rather than a single objective function. EAs reduce the risk of under-sampling, however, they typically require hundreds to thousands of generations with tens to hundreds of individuals — that needs lots of computational resources.

Main advantage of these methods is that they are derivative-free: only the forward pass and objective function is evaluated. This makes them applicable to any model and any metric, including non-differentiable ones. However, the cost is sample efficiency — each candidate requires a full forward simulation, and many candidates are needed per optimization run.

#### 2.2.3 Gradient-based Tuning

Gradient-based optimization requires computing the partial derivative *∂L/∂θ* for all trainable parameters. Doing this analytically is impractical and inefficient for complex simulations with thousands of timesteps. Automatic Differentiation (AD) solves this by decomposing the computation into elementary operations and applying the chain rule systematically. The most common method is reverse-mode AD, also called Backpropagation (BP), is a particularly efficient method: it creates a computational graph of operations — each of them has to be differentiable. Then one backward pass yields all parameter gradients simultaneously. `JAX`, a high performance array computing library, implements this through function tracing and transformation `(jax.grad)`.

A modern instantiation of this approach is `Jaxley` [5], a `JAX`-based simulator that enables gradient-based fitting of HH-type biophysical models at scale(section 3.1.2). However, HH-type models remain computationally expensive at scale, and `Jaxley` does not support simplified models like the AdEx for gradient-based fitting — their discrete reset condition (eq. (3)) breaks automatic differentiation. Addressing this issue requires special treatment: surrogate gradients.

### 2.3 Surrogate Gradients

Surrogate Gradients (SGs) were developed to adapt spiking neurons (usually LIF models) for gradient-based training, primarily on classification tasks in Spiking Neural Networks (SNNs). There, thousands of synaptic weights are trained under a task loss such as cross-entropy, and the surrogate’s role is to let those weight gradients flow through the otherwise non-differentiable spike events. Model parameters are usually fixed (and given as hyperparameters).

To use AdEx for biophysical simulations, the actual model parameters (*g*_*L*_, *a, b, τ*_*w*_, …) — each with different units and scales — need to be learned. Some affect every timestep (leak), others only act at spike events (e.g. reset and adaptation variables). A loss function must quantify similarity between simulated and recorded spike trains. Whether SG, which were shown to work for weight training, also enable biophysical parameter fitting (for simplified neuron models) remains an open question.

Training these models on real data requires differentiating through the forward simulation. Because LIF-type models are discontinuous (see reset in equation 3), automatic differentiation produces undefined or zero gradients. To overcome this issue, prior work replaced the discontinuous (Heaviside-) functions derivative with a (e.g. sigmoid) surrogate, but only for the backward pass. For training weights in SNN, this performed sufficiently well. There exists a variety of common surrogates. They share the commonality that they are centred at the discontinuous point of the Heaviside-function, but differ in shape. To evaluate the differences between surrogate functions, we have to understand how BPTT works.

#### 2.3.1 Backpropagation Through Time

Compared to normal backpropagation, we have a sequence of integration steps *f* (*θ, s*_0_, *t*_0_) *↦ f* (*θ, s*_1_, *t*_1_) *↦* … *↦ f* (*θ, s*_*T*_, *T*), where *θ* are the parameters, *s*_*t*_ is the internal state (voltage *v* and adaptation variable *w*) and *t* describes the time. The loss is defined as *L* = ℒ (*s*_0_, *s*_1_, …, *s*_*T*_). To compute the gradient, we unroll the recurrence into

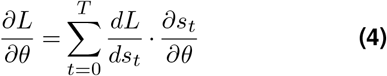

where

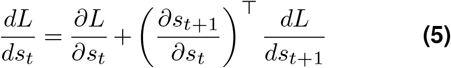

with boundary condition 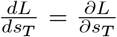, since no future states exist beyond *T*.

Because *s*_*t*_ influences *L* both directly (as an argument to *L*) and indirectly (by affecting *s*_*t*+1_, *s*_*t*+2_, … through the recurrence), we must distinguish between the immediate partial derivative *∂L/∂s*_*t*_ — which considers only the direct dependence — and the total derivative *dL/ds*_*t*_, which accounts for both. Eq. 5 decomposes the total derivative into these two contributions: the direct effect on *L*, and the indirect effect propagated backward from *s*_*t*+1_.

#### 2.3.2 Fundamental Problem in Deep Learning

Sepp Hochreiter first identified the vanishing/exploding gradient problem [8]. The gradient chain in Eq. 4 has the consequence that the error signal at time *t* depends on the product of *T*− *t* Jacobians. If the spectral radius of the matrix is continuously *<* 1, this results in vanishing gradients, for radii *>* 1 the opposite. For the AdEx neuron with leak dynamics (focusing on the voltage component) we obtain:

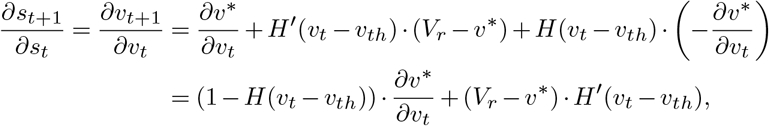

where *H* is the Heaviside-function and *v*^*^ is the Euler integrated voltage (before any reset is applied).

We can infer two conclusions from this equation. First, we can see that *H*^′^ (the derivative of the Heaviside function) is always 0 (or undefined), meaning the reset gradient is ignored. Second, in the subthreshold regime (non-spiking behaviour 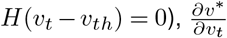 is dominated by the leak current. This leads to vanishing-gradients as visible in Fig. 1.

**Figure 1.**
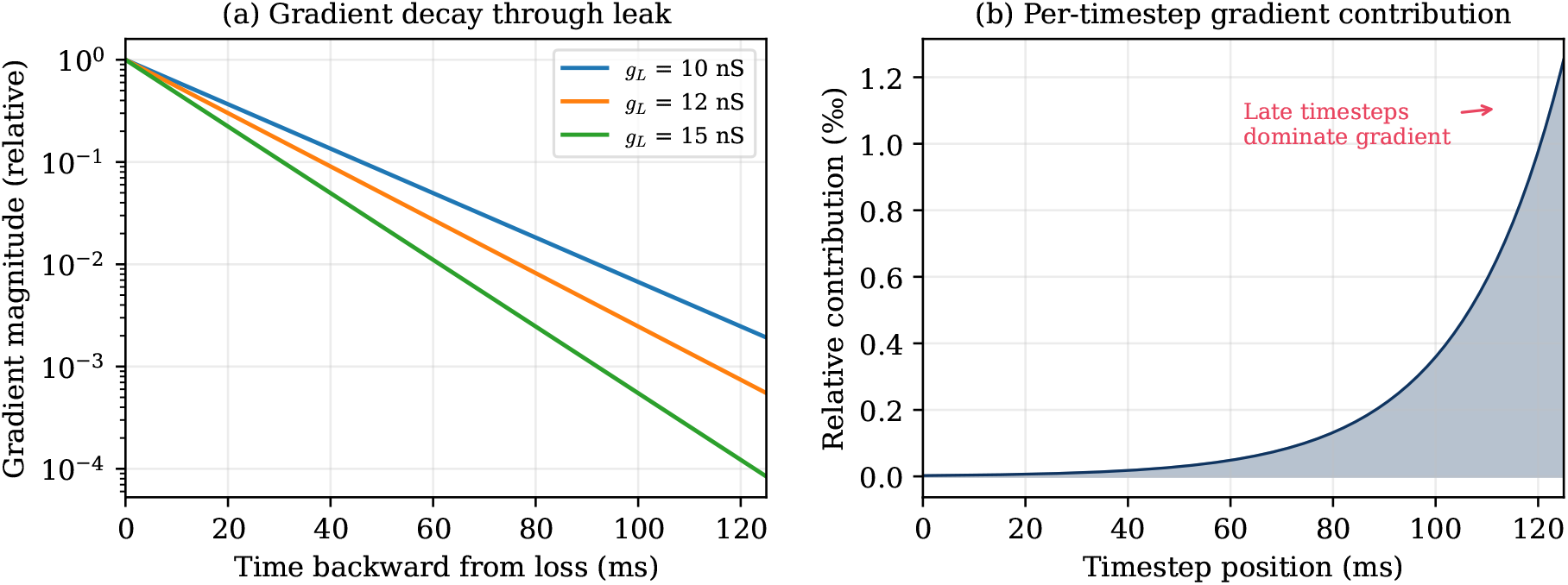
Vanishing gradients through leak dynamics. (a) Gradient magnitude decays exponentially as it propagates backward from the loss, shown for three leak conductance values *g*_*L*_. (b) Consequence for per-timestep gradient contribution: early timesteps contribute negligibly to the total gradient, while the final milliseconds dominate. Computed for *g*_*L*_ = 10 nS, *C* = 200 pF, *dt* = 0.025 ms.

The surrogate gradient approach addresses the first problem by replacing *H*^′^(*v*_*t*_ −*v*_*th*_) with a surrogate derivative *φ*_*β*_(*v*_*t*_ −*v*_*th*_) during the backward pass only, leaving the forward dynamics unchanged:

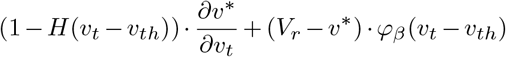

This restores a nonzero gradient pathway through spike events. Crucially, it does not address the second problem — gradient decay through leak dynamics persists — regardless of the surrogate choice.

The severity of this problem differs across parameters. Parameters that affect the continuous subthreshold dynamics (e.g. *g*_*L*_, *E*_*L*_) receive gradient signal at every timestep through the leak term. Parameters that only act through the reset mechanism are much harder to fit. In particular, the spike-triggered adaptation *b* is only updated when a spike occurs (Equation 3: *w* → *w* + *b*). Gradient information for *b* must therefore pass through the Heaviside derivative *H*^′^, which is exactly the term that is zero without surrogates. Even with surrogates, the gradient for *b* is limited to the few timesteps where spikes occur, while the gradient must then survive the leak decay to propagate further backward. Similarly, *τ*_*w*_ controls adaptation dynamics that are most relevant in the post-spike recovery period.

#### 2.3.3 Surrogate gradients

There are a variety of surrogate functions that have been proposed. The simplest one is based on the sigmoid function

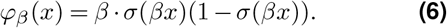

Other common surrogates include the exponential

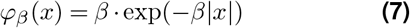

and SuperSpike[9]

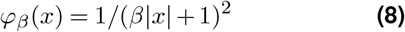

functions. All are centred at the discontinuity and unimodal, but differ in tail behaviour (Figure 2).

**Figure 2.**
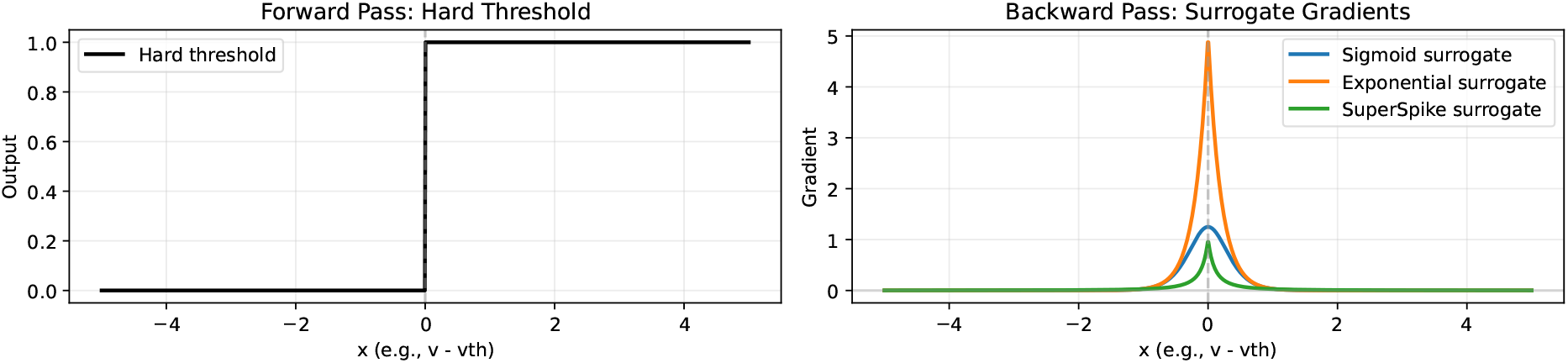
Surrogate gradient shape comparison. (a) The Heaviside step function used in the forward pass, with zero derivative everywhere except at the discontinuity. (b) Three common surrogate derivatives used in the backward pass: sigmoid, exponential, and SuperSpike. All are centred at the threshold and unimodal, but differ in peak height and tail decay. The steepness parameter *β* (here *β* = 5) controls the width of the effective gradient window.

An important hyperparameter is the steepness *β*: large *β* concentrates the gradient signal in a narrow window around threshold, while small *β* spreads it over a wider voltage range. Gygax & Zenke [10] show that surrogate gradients do not need to be the true gradient — it is enough to only point in a useful descent direction.

We will further discuss the choice of surrogate, its steepness and how it interacts with the loss landscape in section 5.2.

### 2.4 Loss Functions for Spiking Models

Surrogate gradients allow gradients to flow backward in time, however, the loss function’s landscape determines the direction of the gradient. If the gradient points towards a non-spiking solution, the model will not produce correct spiking behaviour, no matter the surrogate gradient.

#### 2.4.1 Non-convex Parameter Space

The AdEx parameter space is highly non-convex. Bifurcations in the dynamics — the boundaries between non-spiking, tonic, and bursting regimes characterized by Naud et al. [2] — mean that small parameter changes can cause large qualitative changes in firing behaviour. A typical loss landscape therefore consists of large plateaus, on which the model is silent and the loss is insensitive to most parameters, separated by narrow valleys containing the spike-producing parameter combinations. Gradient descent is consequently sensitive to initialization: a starting point on a non-spiking plateau provides no gradient signal pointing towards the spike-producing region.

#### 2.4.2 Loss function categories

We can differentiate in three category of losses:

1. Pointwise differences
2. Feature-based losses
3. Distance-based metrics

Category one includes Mean Squared Error (MSE), Mean Absolute Error (MAE) and other variations. MSE is defined as

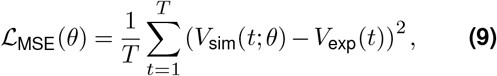

where *θ* denotes the trainable parameters.

Usually these types of losses build a good starting point for lots of machine learning tasks, however, not in this case. Consider two voltage traces with identical spiking frequencies but small temporal misalignments. At each spike in one voltage trace, the other is in subthreshold regime. This leads to high pointwise differences and therefore to high pointwise losses. The crucial problem here is that a non-spiking model would produce a smaller loss than the slightly misaligned one, the gradient therefore pulls the model into non-spiking behaviour (see section 5.2.2, fig. 5).

The second category of loss functions are feature-based losses. Here we evaluate the spike trains for different metrics, e.g. the number of spikes, spiking frequency, time to first spike, mean subthreshold voltage or many more. The simplest one is voltage mean and deviation, but they can be more specific. These loss functions are commonly used for parameter fitting in simplified neuron models. However, by default, most of these features are not in itself differentiable. For non-gradient methods like Grid Search, Nelder– Mead or Genetic Algorithms, this is irrelevant. When applied to gradient-based optimization, the discrete operations inside feature extraction must be replaced by differentiable surrogates. Two primitives suffice in practice. A hard threshold (*did a spike occur?*) is replaced by a sigmoid, and a hard index selection (*which* time step carries the *n*-th spike?) — an argmax over the time axis — is replaced by a temperature-weighted average, i.e. a *soft-argmax*. Writing the per-time-step selection weights as a softmax,

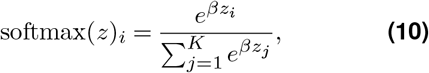

temperature *β* ∈ ℝ^+^, the differentiable feature is for a tuple **z** = (*z*_1_, …, *z*_*K*_) ∈ ℝ^*K*^ and an inverse0020the resulting *expectation ∑* expected spike time), *not* the softmax itself: softmax is a soft argmax, whereas a genuinely smooth max would be the log-sum-exp 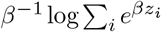. Larger *β* sharpens both towards their hard counterparts. We (e.g. the follow exactly this recipe — sigmoid gates over a cumulative spike count, and a temperature-weighted average of spike times — in section 4.1.2.2. Also important to note is the fact that when different features are combined in a single loss, weighting each features contribution becomes another hyperparameter in the training process.

The third category contains spike-train similarity metrics such as the Van Rossum distance and dynamic time warping. Unlike MSE, these compare spike trains as structured objects rather than sample-by-sample; unlike feature-based losses, they preserve the full temporal structure of the train.

The Van Rossum distance [11] convolves each spike train with a causal exponential kernel *K*(*t*) = exp(−*t/τ*) and computes the *L*^2^ distance between the resulting filtered traces:

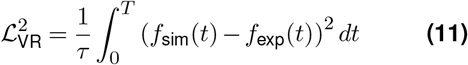

where *f* (*t*) =∑_*i*_ *K*(*t*−*t*_*i*_) 1[*t*≥*t*_*i*_] is the filtered spike train.The time constant *τ* controls the temporal precision: small *τ* penalizes sub-millisecond timing errors (temporal coding), while large *τ* smooths over individual spikes and compares overall firing rates (rate coding). Because convolution and the *L*^2^ norm are smooth operations, the Van Rossum distance is naturally differentiable — no special treatment is required for the loss itself.

An alternative approach is Dynamic Time Warping (DTW) [12], which finds an optimal alignment between two time series by warping the time axis. Given two sequences *x* ∈ ℝ^*n*^ and *y* ∈ ℝ^*m*^ with pointwise cost matrix *δ*_*i,j*_ = (*x*_*i*_ − *y*_*j*_)^2^, the DTW distance is defined via the recursion

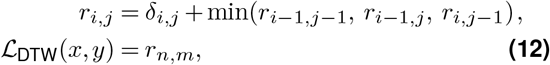

with boundary conditions *r*_0,0_ = 0 and *r*_*i*,0_ = *r*_0,*j*_ = ∞for *i, j >* 0. Unlike pointwise losses, DTW tolerates temporal shifts by design; however, the hard minimum in Eq. **(12)** is not differentiable. Soft-DTW [13] replaces it with the softmin

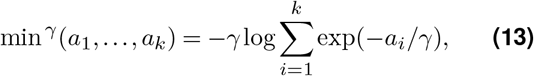

yielding the smoothed recursion

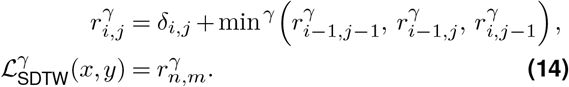

The smoothing factor *γ >* 0 controls the relaxation: as *γ*→ 0, Soft-DTW recovers the original DTW distance, while larger *γ* produces smoother gradients at the cost of a looser alignment.^*^ This makes the alignment distance amenable to gradient-based optimization, at the cost of *O*(*nm*) time and memory complexity.

All three categories share one fundamental constraint: every operation in the computational path from parameters to loss must be differentiable. For pointwise losses, this is trivially satisfied. For feature-based losses, operations like spike detection and counting must be replaced with soft, differentiable approximations. For distance metrics operating on spike trains, the spike train itself must be represented in a differentiable form. This requirement ties directly back to the surrogate gradient framework introduced in Section 2.3: the surrogate provides a soft spike indicator *s*(*t*) ∈ [0, 1] that can serve as input to any of these loss functions.

### 2.5 Summary

The AdEx model provides an efficient yet expressive neuron model, but parameters must be fitted through optimization to get expressive, neuron imitating firing behaviour. Gradient-based methods are attractive due to their computational efficiency, however, the Heaviside reset mechanism creates a differentiability barrier that prevents standard automatic differentiation. Surrogate gradients restore gradient flow through spike events by replacing the Heaviside derivative with a smooth surrogate during the backward pass. Three challenges remain: the parameter space is highly non-convex, gradients still decay exponentially through the leak dynamics, and the surrogate steepness *β* must be chosen carefully. The loss function is equally critical. MSE on raw voltage traces drives the optimizer towards non-spiking solutions, feature-based losses lack temporal credit assignment, and distance metrics like Van Rossum or soft-DTW offer promising alternatives but come with their own trade-offs. The interplay between surrogate choice and loss function design is the central challenge investigated in this work.

## 3 Related Work

Simplified neuron models were originally a necessity: computing resources were too limited for large-scale simulations with detailed ion-channel models. As hardware improved, focus shifted to biophysically detailed HH-type models, most notably through the Blue Brain Project [14] and the Human Brain Project[15]. In recent years, however, interest in simplified models resurged, driven by two developments: the use of spiking neurons in machine learning [16] and the ambition to simulate brain-scale networks where computational efficiency remains essential [3].

### 3.1 Parameter Estimation for Neuron Models

#### 3.1.1 Derivative-free Methods

Simplified neuron models like the AdEx contain parameters (e.g. *a, b, τ*_*w*_) that lack direct physiological counterparts and must therefore be estimated through optimization. Van Geit et al. [6] survey the landscape of automated parameter search methods for neuron models, ranging from evolutionary strategies to simulated annealing.

BluePyOpt [14] operationalized multi-objective evolutionary optimization for neuroscience, becoming the most widely used tool for automated parameter fitting. It wraps the DEAP library, providing access to IBEA, NSGA-II, CMA-ES and particle swarm optimization, and computes fitness using the eFEL (Electrophysiology Feature Extraction Library) to compare features extracted from simulated and recorded voltage traces. BluePyOpt treats each feature as a separate objective, enabling Pareto-front-based selection without requiring a single scalar loss function. However, being population-based, it requires hundreds to thousands of forward simulations per optimization run.

For the AdEx model specifically, Marín et al. [17] used genetic algorithms and Cruz et al. [18] applied Teaching-Learning-Based Optimization (TLBO) and multi-state single-agent stochastic search (MSASS) to fit AdEx parameters to voltage-clamp recordings, demonstrating that derivative-free methods can successfully recover AdEx parameters. Guarino et al. [19] took a different approach: they extracted eight electrophysiological features (time to first, second, third and last spike, inverse of first and last Inter-Spike Interval (ISI), firing frequency, and voltage at stimulus end) and performed parameter space exploration using Sobol quasi-random sequences to sample candidate models. Each model is ranked by the sum of relative errors across all features, with a penalty of +3 for missing features (e.g. when a non-spiking model cannot produce spike times). Their work demonstrates that feature-based fitting can succeed for the AdEx model across seven neuron types and five brain regions, but relies on brute-force search rather than gradient-based optimization and does not evaluate using the coincidence factor Γ.

#### 3.1.2 Gradient-based Methods

Using gradient descent for parameter fitting requires that the entire computational path — from parameters through simulation to loss — is differentiable. For HH-type models, this is inherently the case: the ion channel dynamics are described by smooth differential equations with no discontinuities. Jones et al. [7] demonstrated that gradient-based methods can efficiently optimize ODE-based neuron models using automatic differentiation. Hertäg et al. [20] developed an analytical approximation to the AdEx model to speed up computations, though this sacrifices the model’s full expressive power.

Deistler et al. [5] developed `Jaxley`, a `JAX`-based simulator that enables gradient descent for biophysical models at scale. They fitted single-neuron models with up to 1,390 parameters and trained networks with 100,000 parameters. On synthetic data, gradient descent required only ~9 optimization steps (median across runs) to find parameter sets matching the ground truth, while the IBEA genetic algorithm needed a similar number of generations with ten simulations each — making gradient descent roughly an order of magnitude more sample-efficient. On experimental recordings from the Allen Cell Types Database, 1,000 parallelized gradient descent runs on a GPU produced fits that closely matched recorded voltage traces. However, `Jaxley` operates exclusively on HH-type models. Simplified models with discrete reset mechanisms (Section 2.3) break the differentiability assumption and are not supported for gradient-based fitting.

#### 3.1.3 Simulation-based Inference

A third paradigm avoids optimization entirely: simulation-based inference (SBI) uses neural density estimators to approximate the posterior distribution *p*(*θ* |**x**) over parameters given observed data. Gonçalves et al. [21] developed Sequential Neural Posterior Estimation (SNPE), which trains a deep neural network on simulated data to learn the mapping from observations (or summary statistics) to the full posterior. Unlike point-estimate methods, SNPE captures the complete space of data-consistent parameters, including parameter degeneracies and multimodal solutions. It is amortized: after an expensive initial training phase, inference on new recordings is nearly instantaneous. Critically, SNPE treats the simulator as a black box and can therefore be applied to non-differentiable models including simplified spiking neurons. It comes from the same research group as Jaxley (Macke lab) and represents the Bayesian counterpart to the gradient-based approach.

### 3.2 Surrogate Gradients in Spiking Neural Networks

Surrogate gradients were developed to enable gradient-based training in SNNs, where the discrete spike mechanism prevents standard backpropagation. Neftci et al. [16] provide a comprehensive tutorial on the approach, covering sigmoid, exponential and piecewise-linear surrogates for training LIF-based networks on classification tasks. Gerum and Schilling [22] systematically compared surrogates (sigmoid, Esser function, and others) when training LIF neurons integrated into standard machine learning architectures via Keras, finding that classification performance was largely robust to the specific surrogate choice. Similarly, Perez-Nieves et al. [23] used a sigmoidal surrogate in `PyTorch` to train spiking networks on classification benchmarks, demonstrating that heterogeneous neuron parameters improve learning — though these parameters were drawn from distributions, not fitted to data.

Zenke and Ganguli [9] proposed SuperSpike, a surrogate based on 1*/*(*β* |*x*| + 1)^2^, and formally analysed the vanishing gradient problem in multi-layer SNNs. They showed that careful surrogate design is essential for learning in deep spiking networks, particularly for precise spike timing tasks.

Recent theoretical work by Gygax and Zenke [10] provides a formal foundation for why surrogate gradients work despite not being the true gradient. They show that, for single neurons, surrogate gradient descent is equivalent to computing derivatives of the expected output in smoothed probabilistic models — the surrogate derivative matches the derivative of the neuronal escape noise function. This equivalence breaks down in multilayer networks, where surrogate gradients remain a heuristic. Importantly, SNN training has been shown to be robust to the choice of surrogate function [10], suggesting that the exact surrogate shape matters less than providing *any* non-zero gradient through spike events.

All of the above work focuses on training *synaptic weights* for task-based objectives (classification accuracy, spike timing). Whether surrogate gradients also enable fitting *biophysical parameters* (*g*_*L*_, *a, b, τ*_*w*_) to match experimental recordings remains unexplored. This is a fundamentally different problem: the parameters are heterogeneous (different units, scales, and roles), the loss must quantify spike train similarity rather than task performance, and the surrogate must provide useful gradient information for parameters that interact with the spike mechanism in qualitatively different ways (Section 2.3).

A complementary line of work avoids surrogates altogether by exploiting neuron models in which spike times depend smoothly on parameters. Klos and Memmesheimer [24] show that, for the Quadratic Integrate-and-Fire (QIF) neuron, the infinite voltage slope at threshold guarantees that small parameter changes shift spike times continuously rather than causing them to appear or disappear mid-trial. Combined with their *pseudospike* construction — an extension of the dynamics that keeps a “would-be” spike time addressable past the trial end, even for currently silent neurons — this yields exact, event-based gradients without any surrogate, and resolves the dead-neuron problem by construction. The approach is restricted to neuron models with closed-form between-spike solutions and infinite spike-initiation slope, which excludes the AdEx model considered in this thesis but provides a useful theoretical contrast: surrogate gradients trade exactness for generality across neuron models.

### 3.3 Benchmarking Neuron Models

Comparing the quality of neuron model fits across studies requires a standardized evaluation metric. Jolivet et al. [25] introduced the coincidence factor Γ as a measurement of the fraction of coincident spikes between a model and a reference recording within a temporal precision window Δ (2 ms in the original paper), corrected for chance:

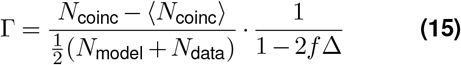

where *N*_coinc_ is the number of coincident spikes within ±Δ, *⟨N*_coinc_*⟩* is the expected number of coincidences under a Poisson model with the same firing rate (chance level), and *f* is the model firing rate. The factor 1 −2*f* Δ normalizes Γ such that Γ = 1 corresponds to a perfect match and Γ = 0 to chance-level performance.

In their benchmark, Jolivet et al. evaluated several simplified neuron models fitted using the Nelder–Mead simplex algorithm. The best-performing models — including the AdEx — achieved Γ ≈0.82–0.83 on heldout test data [25], establishing a concrete performance target for neuron model fitting. These results demonstrate that simplified models *can* reproduce experimental spike trains with high fidelity when their parameters are fitted appropriately.

The precision parameter Δ in Γ controls the temporal tolerance for counting coincident spikes: small Δ requires near-exact spike alignment, while large Δ tolerates timing errors. This mirrors the role of the time constant *τ* in the Van Rossum distance (Section 2.4.2), where small *τ* emphasizes temporal coding and large *τ* emphasizes rate coding. In both metrics, a tolerance parameter (Δ or *τ*) sets how strictly spike times must align to count as a match.

### 3.4 Loss Functions for Neuron Model Fitting

The choice of loss function is critical for any optimization approach, but becomes especially consequential for gradient-based methods, where the loss determines not only the optimization target but also the gradient direction.

#### 3.4.1 Feature-Based Losses

Guarino et al. [19] define a loss based on the relative error between data-derived and model-derived electrophysiological features (the eight features listed in section 3.1), with a fixed penalty for features that are present in the data but absent from the simulation (e.g. a non-spiking model cannot produce spike times). This formulation works well with derivative-free search, but the features themselves are not differentiable: spike detection requires threshold crossings, and spike counting is discrete. Adapting these features for gradient-based optimization requires differentiable approximations (soft threshold, soft spike count), which we investigate in this work.

#### 3.4.2 Summary Statistics and Soft-DTW

Deistler et al. [5] use two complementary loss functions for fitting HH-type models. For synthetic experiments, they define four differentiable summary statistics: the mean and standard deviation of the voltage trace in two time windows (pre-stimulus and stimulus), normalized by fixed constants (8.0 and 4.0 respectively). The loss is the MAE on these standardized statistics. This suffices for coarse parameter recovery, where the goal is matching the overall voltage distribution rather than individual spikes.

For fitting to experimental recordings, they switch to soft-DTW [13], which tolerates temporal misalignments by warping the time axis. To make DTW practical for long traces, they preprocess by applying a sliding window maximum (window size 50, stride 30 timesteps at Δ*t* = 0.025 ms) and rescaling both signals to the unit interval. Critically, they augment the cost matrix with a temporal penalty, yielding the cost function *c*(*x*_*i*_, *y*_*j*_) = |*x*_*i*_ −*y*_*j*_ |+ |*i*− *j*|, which prevents pathological warping paths. This approach produced good fits on Allen Cell Types Database recordings, but was applied exclusively to HH-type models where no surrogate gradients are needed.

#### 3.4.3 Spike Train Distance Metrics

The Van Rossum distance [11] (eq. (11)) is naturally differentiable and therefore a candidate for gradient-based optimization, but it has primarily been used as an *evaluation* metric in theoretical neuroscience rather than as an *optimization* objective. The related Victor–Purpura metric is based on edit distances (spike insertion, deletion and shifting) and is not differentiable.

Existing work falls into two non-overlapping categories: derivative-free methods applied to simplified models using any loss function [17–19], and gradient-based methods applied to inherently differentiable HH-type models [5, 7]. No prior work systematically investigates loss function design for gradient-based optimization of simplified neuron models that require surrogate gradients.

### 3.5 Training Stabilization for Biophysical Models

Training biophysical models with gradient descent is challenging beyond the choice of loss function. Gradient norms can spike by orders of magnitude during action potentials, parameters have different scales and units, and the loss landscape is highly non-convex with many local minima [5].

Deistler et al. [5] describe several stabilization techniques implemented in `Jaxley`. First, they reparameterize bounded parameters through an inverse sigmoid transform that maps *θ* ∈ [*l, u*] to an unconstrained variable, simultaneously placing all parameters on a comparable scale and eliminating the gradient discontinuities at the bounds that hard clipping would introduce. Second, they employ Polyak stochastic gradient descent, in which the update is normalized by the gradient norm and rescaled by the current loss value, yielding implicit learning-rate decay as the optimum is approached. Third, they assign different optimizers to different parameter groups (e.g. ion channel conductances vs. morphological parameters), since gradient norms differ systematically across parameter types. Fourth, they use multilevel checkpointing to reduce the memory cost of backpropagation through long simulations, and in some experiments truncate the gradient in time to mitigate vanishing or exploding gradients.

To address the non-convex landscape, Deistler et al. parallelize up to 1,000 gradient descent runs from different initializations on a single GPU, selecting the best parameter set post-hoc. This multi-start strategy is effective because individual runs are cheap once the simulation is compiled.

These techniques are available as building blocks in `Jaxley` but have only been demonstrated for HH-type models, where gradients flow naturally through smooth ion channel dynamics. For simplified models with surrogate gradients, additional challenges arise: the surrogate introduces an approximation error in the gradient, adaptation parameters (*b, τ*_*w*_) receive gradient signal only through spike events, and the effective gradient window of the surrogate may not match the voltage range where parameters are most sensitive. Whether the same stabilization recipe transfers to this setting is an open question that we investigate empirically.

### 3.6 Summary and Research Gap

The landscape of neuron model parameter fitting spans three paradigms. Derivative-free methods (EAs, Nelder–Mead, parameter space exploration) have been successfully applied to the AdEx model [17–19], but require many forward simulations and scale poorly to large parameter counts. Gradient-based methods achieve orders-of-magnitude speedups for HH-type models [5, 7], but rely on the inherent differentiability of ion channel dynamics. Simulation-based inference [21] provides full posterior distributions without differentiability requirements, but targets a different goal (uncertainty quantification) and requires a separate training phase.

Surrogate gradients bridge the differentiability gap for simplified models, enabling gradient flow through discrete reset events. However, prior work has validated surrogates exclusively for training synaptic weights on task-based classification objectives [9, 16, 22, 23]. Fitting biophysical parameters to experimental recordings poses distinct challenges: heterogeneous parameters, spike train similarity losses, and vanishing gradients through leak dynamics.

No prior work systematically investigates how loss function design interacts with surrogate gradient optimization for simplified neuron models. This thesis addresses that gap.

## 4 Methods

This chapter describes how we fit the parameters of the AdEx model and how we measure whether a fit has succeeded. Designing a loss whose surface a gradient can usefully descend is itself part of the research problem here, not a preliminary to it: we develop and compare three differentiable loss functions MSE, the feature-based loss of Guarino et al., and the Van Rossum distance. Around them we set out the evaluation metric and recovery protocol, the training-stabilization applied on top of the optimizer, and two benchmarks — a recovery-radius benchmark against derivative-free baselines (Nelder–Mead) and a spike-free subthreshold benchmark that isolates gradient flow from the reset. The chapter closes with the implementation these experiments rely on (section 4.3): the differentiable AdEx model in `Jaxley` and its surrogate-gradient variant, both verified against Brian2.

### 4.1 Parameter Optimization

To evaluate the effectiveness of gradient-based optimization using surrogate gradients, we recover parameter of synthetic voltage traces. This way, the ground truth is known, so recovery success is measurable. We perform a perturbation protocol, meaning we identify in which distance from the ground truth the optimization setup can reliably recover the parameters.

The full protocol — scenarios, perturbation grid, and the four methods we compare — is specified in section 4.1.4.

#### 4.1.1 Evaluation Metrics

Deistler et al. [5] claim sub-millisecond and sub-millivolt matching for their HH-type fits; for simplified models against real experimental data this is an unrealistic expectation, and the natural goal is to replicate spike timings as well as possible. We therefore evaluate every fit with the coincidence factor Γ defined in eq. (15) (section 3.3), using the conventional precision window Δ = 2 ms. We clip Γ to [ − 1, 1]: large negative values arise from non-spiking models and signal only that the optimizer failed to escape its initial state, with no further information beyond that fact. Because our recovery experiments use synthetic targets generated by the same model family, full recovery yields Γ = 1; this is a stricter bar than Jolivet et al.’s Γ ≈ 0.82– 0.83 on experimental data (section 3.3), which reflects the model–data mismatch that is absent in our setup. We count a recovery as successful when Γ *>* 0.5. Since Γ is normalized so that Γ = 0 corresponds to chance-level overlap (a Poisson process of the same rate) and Γ = 1 to perfect coincidence, Γ *>* 0.5 identifies fits whose spike timing agrees with the target more than it disagrees relative to chance. It is the conventional midpoint used in the coincidence-factor literature, and because Γ increases monotonically with timing agreement the precise cut-off is not critical: 0.5 cleanly separates fits that have captured the target’s firing pattern from those that have not.

#### 4.1.2 Loss Functions

We compare three differentiable losses on the synthetic-recovery benchmark, each chosen for a specific role: MSE as a baseline that exposes the timing-agnostic failure mode of pointwise losses (section 2.4.2); the feature-based loss of Guarino et al. [19] to study what softening discrete features for gradient descent costs in practice; and the Van Rossum distance as a spike-train metric whose kernel time constant *τ* exposes the precision/rate-coding trade-off through a single hyperparameter. The remainder of this subsection fixes the implementation details that are specific to our discrete-time, surrogate-gradient setting; the underlying definitions are recalled by reference to section 2.4.2.

##### 4.1.2.1 MSE Loss

We include the MSE loss defined in eq. (9) not as a candidate loss for parameter fitting, but as a baseline against which the timing-aware losses are compared. Its role is to empirically confirm the failure mode argued on theoretical grounds in section 2.4.2: that a pointwise loss on raw voltage drives the optimizer towards non-spiking solutions whenever the simulated and target spikes are not perfectly aligned.

Concretely, the loss is evaluated between the simulated voltage trace *V*_sim_(*t*; *θ*) produced by the AdExSurrogate channel and the target trace *V*_exp_(*t*), sampled at the simulation timestep Δ*t* over the full stimulus window [0, *T*]. No masking, windowing, or normalization is applied; both traces enter the loss in millivolts.

In contrast to the feature-based and Van Rossum losses discussed below, MSE requires no soft approximations on the loss side — the squared difference and the temporal mean are smooth operations. The only non-differentiable element on the path from parameters to loss is the spike reset of the AdEx model itself, which is handled by the surrogate gradient described in section 4.3.2.1. This makes MSE the cleanest probe of the surrogate gradient in isolation, uncontaminated by the additional softness hyperparameters that the other losses introduce. For the same reason, MSE has no tunable hyperparameters of its own: any observed failure cannot be attributed to a poor choice of softness, weighting, or kernel parameters, and must instead reflect a property of the loss itself. The corresponding training results are presented in section 5.2.2 and fig. 5.

##### 4.1.2.2 Feature-Based Loss

Simplified neuron models are usually not required to perfectly imitate the voltage trace of a neuron — their main requirement is to faithfully replicate the firing behaviour. A common approach is therefore to extract electrophysiological features from the experimental and simulated traces and compare these instead, e.g. via the eFEL library [26]. We adopt the eight features identified by Guarino et al. [19] (section 3.1): four spike-time features (*t*_1_, *t*_2_, *t*_3_, *t*_last_), two ISI-inverse features (1*/*ISI_1_, 1*/*ISI_last_), the mean firing frequency, and the membrane voltage at stimulus end (*V*_stimend_).

The loss is computed as a weighted sum of relative errors:

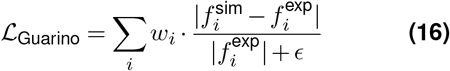

where 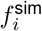 and 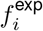 denote simulated and experimental feature values respectively, and *w*_*i*_ are feature-specific weights. When a feature is present in the experimental data but absent in the simulation (e.g., the experimental trace has three spikes but the simulation produces only one), a fixed penalty is added to encourage the optimizer to produce the correct number of spikes.

This feature-based formulation addresses the spike timing sensitivity problem inherent to MSE: small temporal misalignments no longer produce catastrophic loss values, as long as the overall firing pattern — captured by the features — remains similar.

The feature-based loss requires extracting spike times and counts from voltage traces. Standard implementations use hard threshold crossings, which are non-differentiable. To enable gradient-based optimization, we developed soft approximations of the feature extraction operations.

The naming “soft approximation” is potentially misleading and warrants up-front clarification: the spike trace itself is never softened. The `AdExSurrogate` channel emits a strictly binary indicator *s*(*t*) ∈ {0, 1} at every time step (section 4.3.2.1), so the recorded trace contains real, mass-1 spike events^*^ — there are no fractional spikes anywhere in the forward pass. In particular, the spike count *N*_spikes_ =*s*(*t*) and the cumulative count ∑_*τ*≤*t*_ *s*(*τ*) are exact integers.

What is “soft” is the *recipe* we use to turn this binary signal into feature values, not the values it produces. Concretely, the time of the *n*-th spike could in principle be read off as argmin *t* : ∑_*τ* ≤*t*_ *s* (*τ*) = *n*} Δ*t*, but argmin is non-differentiable: a small change in *θ* leaves the integer index unchanged (then jumps it by one), so the chain-rule derivative through such an operation is zero almost everywhere. We replace this hard lookup with a smooth weighted average over time, gated by a sigmoid-of-cumulative-sum selector (described in detail below). On a binary *s*(*t*) the selector collapses to ≈ 1 at the *n*-th spike’s index and ≈0 elsewhere, so the forward value is essentially identical to the exact spike time; what differs is that the recipe is now a chain of differentiable operations. The temperature *τ* and sigmoid sharpness *β* control how sharp this selector is; they do not turn *s*(*t*) itself into a continuous variable.

Validity flags (sigmoid gates of the form 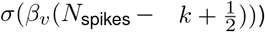 follow the same pattern: they are sigmoid-smoothed indicators of integer comparisons, exact in forward but with a non-zero analytical derivative.

Differentiability with respect to the model parameters is supplied at exactly one node, the `mcustom_vjp` on *s*(*t*): although *s* is a step function with true derivative zero almost everywhere, section 4.3.2 substitutes the smooth surrogate kernel *∂s/∂v* during the backward pass. The chain rule then propagates a finite *∂ℒ /∂θ* through the smooth feature recipe and back to the parameters. The resulting gradient does not correspond to any finite-difference test of the loss — the true loss landscape is piecewise constant in *θ* and finite differences would return zero. It is a surrogate gradient in the sense of Neftci et al. [16]: a useful optimization direction, formally interpretable as the exact gradient of a smoothed version of the model, but not the true derivative of the staircase loss. A consequence of keeping *s*(*t*) binary is that the Van Rossum distance (section 4.1.2.3), which consumes *s*(*t*) directly, sees mass-1 events on both the simulated and target side, with no per-event mass mismatch.

For spike timing, we use cumulative sums to identify when the *n*-th spike occurs and compute a weighted average over time. Similarly, ISI features are derived from the difference between consecutive soft spike times. Features that require a minimum number of spikes (e.g., ISI needs at least two spikes) are weighted by soft validity flags that smoothly transition based on the spike count.

The soft approximations introduce two hyperparameters: a temperature *τ* that controls how sharp the spike selection is, and a sigmoid steepness *β* for the masking operations. We used *τ* = 0.3 and *β* = 10.0 based on hyperparameter sweeps.

We note that this differentiable feature extraction is a working implementation that has not been thoroughly validated. Due to time constraints, we did not perform extensive testing of the soft approximations against their hard counterparts. The approach produces gradients and enables optimization, but the accuracy of the soft features compared to exact values remains to be studied in future work.

##### 4.1.2.3 Van Rossum Distance

The definition of the Van Rossum distance is recalled in eq. (11); this subsection only fixes the input we feed it and the implementation choices that follow from our discrete-time, surrogate-gradient setting.

The two spike trains in eq. (11) are the binary indicator *s*(*t*) produced by the `AdExSurrogate` channel for the simulation, and the binary spike train obtained from the experimental voltage by rising-edge threshold crossing for the target. As established in section 4.1.2.2, both are mass-1-per-spike sequences, and gradient flow into the parameters is supplied by the `custom_vjp` on *s*(*t*) rather than by any softening on the loss side. Van Rossum is the only loss in our comparison where this is the case — the convolution with *K* and the *L*^2^ norm are themselves smooth, so no temperature or softness hyperparameter is required to make the path from *s*(*t*) to the loss differentiable.

We use the *squared* Van Rossum distance as the loss (following the 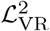 convention of eq. (11) and van Rossum [11]; the square avoids a 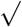 whose only effect is a monotone rescale with a gradient singularity at zero), discretizing eq. (11) on the simulation grid:

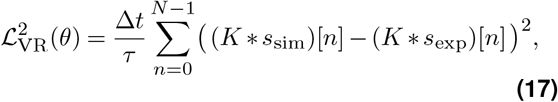

with *K*[*n*] = exp(−*n* Δ*t/τ*) truncated at 5*τ* (where *K*(5*τ*) ≈ 0.007). Truncation keeps the convolution *O*(*Nτ/*Δ*t*) per evaluation rather than *O*(*N*^*2*^); it is exact to the displayed precision in all results we report.

To keep the loss responsive in regimes where the simulated cell produces no spikes at all — in which case eq. (17) reduces to a constant determined entirely by the target and provides no parameter-dependent signal — we optionally add a subthreshold voltage MAE term:

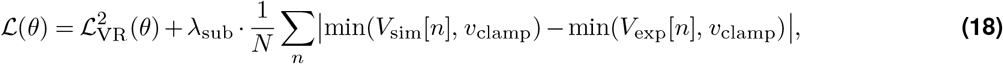

clamped above *v*_clamp_ = −40 mV so that spike peaks contribute nothing and the term reflects only the subthreshold trajectory. *λ*_sub_ is set to 0 when we want to study the spike-train objective in isolation, and to a small positive value otherwise. The role of this term is identical to that of the spike-count regularizer in the feature-based loss: provide a non-zero gradient on the no-spike plateau so the optimizer is not stuck there before the spike-train term has anything to act on.

##### 4.1.2.4 Soft-DTW and Summary Statistics

For completeness, we also implemented two further differentiable losses from the HH-fitting recipe of Deistler et al. [5]: Soft Dynamic Time Warping (SDTW) [13] with their temporally-augmented cost *c*(*x*_*i*_, *y*_*j*_) = | *x*_*i*_ − *y*_*j*_| + |*i*− *j*|, and the four-statistic MAE loss (mean and standard deviation of the voltage trace in the pre-stimulus and stimulus windows) they use for synthetic experiments. Neither produced parameter recoveries comparable to the feature-based or spike-train losses on the AdEx recovery benchmark. We exclude both from the method comparison of section 4.1.4. SDTW is nonetheless retained in the loss-landscape characterization of section 5.2 and the appendix landscape grids, where its narrow, deep global-minimum valleys indicate its inability to recover any but the smallest perturbations. The four-statistic summary loss is not analysed further.

#### 4.1.3 Training Stabilization

Even with surrogate gradients in place and a differentiable loss, training the AdEx model is numerically delicate. Three properties of the problem make plain stochastic gradient descent unreliable: gradient magnitudes vary by several orders of magnitude across parameters because the AdEx parameters live on physically different scales (capacitance in pF, time constants in ms, conductances in nS); the loss landscape contains large non-spiking plateaus where soft spike-detection terms dominate the gradient signal (section 5.2); and the trainable parameters are bounded by physiologically plausible ranges, so naive updates can step outside the feasible region. We address these issues with a small set of stabilization techniques applied on top of the optimizer, summarized below and combined as listed in table S1.

##### 4.1.3.1 Polyak step-size normalization

Polyak’s adaptive step size [27] replaces the fixed learning rate *η* with one that depends on the current loss value and gradient norm:

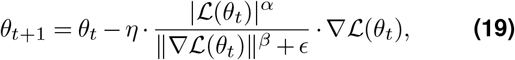

with *α, β* ∈ [0, 1] controlling how strongly the loss and the gradient norm enter. For *α* = *β* = 1 this reduces to a normalized gradient step whose magnitude is set by the current loss, so the update size shrinks automatically as the optimizer approaches a minimum and grows when far from it.

In our setting Polyak provides two practical benefits: with *α >* 0 step sizes decrease with decreasing loss |ℒ|. The *β* variable controls how much of the gradient norm is divided out of the update: *β* = 0 leaves spikes intact (vanilla magnitude), *β* = 1 fully decouples step size from |*∇*ℒ|. The first benefit being automatic annealing when we get close to the global minimum while simultaneously soft capping gradient spikes. Note that this does not address magnitude differences between gradient parameters, because all partial differentials are normalized by the same factors. Large gradient spikes in one parameter actually dampen small gradients of other parameters. Polyak addresses *temporal* gradient smoothing, not *spatial*.

##### 4.1.3.2 Sigmoid parameter transformation

Hard projection of *θ*_*t*_ back into [*θ*_min_, *θ*_max_] after each update introduces two problems for adaptive optimizers: the projected portion of the step is discarded while the optimizer’s internal state (momentum, second-moment estimates) is updated as if the full step had been taken, causing state-trajectory mismatch; and in multi-parameter updates the projection alters the descent direction in a way that depends on the bounds rather than on the loss. Following the transform pattern provided by `Jaxley` [5], we instead reparameterize each bounded parameter through a sigmoid,

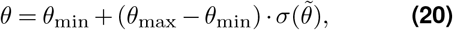

and optimize in the unconstrained space 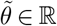. Bounds are then enforced implicitly — *θ* can never leave [*θ*_min_, *θ*_max_] — and the gradient with respect to 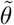 remains finite everywhere, including at the boundary, so the optimizer keeps receiving signal as *θ* approaches a bound rather than having it discarded by a projection step. A side effect is that gradients are smallest near the bounds, which acts as an implicit bias towards central values within each interval. However, this introduces a soft bias and hidden hyperparameters (*θ*_min_, *θ*_max_) for each parameter. Changing these can drastically affect convergence performance. Sensitivity to bound choices is common in bounded optimization, but the sigmoid amplifies it.

##### 4.1.3.3 Multi-trace optimization

Single-trace fitting (especially on experimental data) is underdetermined: many parameter combinations produce the same firing pattern under one stimulus protocol but disagree under others. To exploit this, we extend the loss to a sum over *K* stimulus protocols sharing the same parameters,

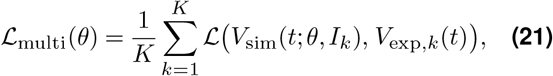

where {*I*_*k*_} is a fixed set of step currents at different amplitudes. The averaged gradient combines temporal credit assignment from regimes that elicit spikes with subthreshold information from regimes that do not, constraining adaptation parameters (*a, b, τ*_*w*_) that are otherwise weakly identifiable from a single trace.

Multi-trace optimization was implemented but is not used in the recovery benchmark of section 4.1.4. Two reasons: it multiplies simulation cost by *K* at every gradient step, which made the per-trial budget incompatible with the sweep size we wanted to run; and the averaged loss is fragile when one of the *K* stimulation regimes drives the model into a non-spiking or numerically chaotic regime, in which case a single trace can dominate *∇*ℒ_multi_ and overwhelm signal from the others. Both issues are addressable in theory: trace selection or per-trace loss weighting for the second, vmap-batched simulation for the first. Neither was within the scope of this thesis. We expect this technique to be necessary rather than optional once the target is experimental data, where ground-truth parameters are not available to confirm recovery and identifiability must come entirely from the loss.

##### 4.1.3.4 Standard regularizers

We also evaluated the more common stabilizers — gradient-norm clipping and exponential learning-rate decay — as drop-in alternatives to Polyak when the latter is disabled. These were ultimately not retained in the reported configurations (table S1) because, on the diagnoses summarized in section 5.2 and section 6, clipping and decay only modulate the magnitude of the update and cannot recover information that is already lost on the non-spiking plateau.

#### 4.1.4 Recovery-Radius Benchmark

The training notebooks (section 5.3.1 and the per-loss subsections of section 5.2) establish that gradient-based optimization with surrogate gradients and Polyak step-size normalization *can* recover *some* AdEx membrane parameters in some configurations. What they do not establish is *how reliable* the recovery is, or how it compares against derivative-free baselines on the same target. A worked example shows what the optimizer does at one operating point; to draw conclusions about gradient-based fitting as a method, we need a protocol that varies the difficulty of the recovery task systematically and that compares methods on matched problem instances. This subsection specifies that protocol; the corresponding results are reported in section 5.3.2.

##### 4.1.4.1 Protocol

We want to observe recovery behaviour in different spiking regimes. For that reason, we used three scenarios: tonic, adaptation and initial bursting — each corresponding to one of the Naud reference parameter sets [2]. We treat the ground-truth parameter vector *θ*^*^∈ ℝ^|*θ*|^ as the recovery target and generate a target voltage trace by simulating the validated AdEx channel of section 4.3.1 under a fixed step current. Then, we construct a family of perturbed initializations

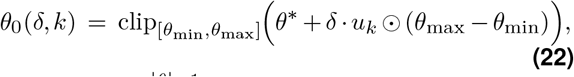

where *u*_*k*_ ∈ S^|*θ*|−1^ are unit vectors drawn uniformly on the sphere (Gaussian sampling followed by *L*^2^-normalization), *δ* is a relative perturbation scale, and the elementwise product ⊙ with (*θ*_max_ −*θ*_min_) rescales each parameter to its bound width so that *δ* has the same meaning across heterogeneous units (pF, nS, mV, ms). The clip operation handles the case where a perturbed component falls outside its parameter bounds.

For every (scenario, *δ, k*) we run each method from *θ*_0_(*δ, k*) to obtain a recovered parameter vector 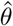, and report the coincidence factor 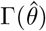 (eq. (15), Δ = 2 ms) computed on a clean (non-surrogate) forward simulation at 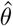. A run counts as *successful* when 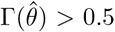, following the success threshold fixed in section 4.1.1. The headline statistic is the success probability *P* (Γ *>*0.5 | *δ*) as a function of the perturbation scale, faceted by scenario and one curve per method; the *recovery radius* of a method is then the largest *δ* for which *P* (Γ *>* 0.5) ≥ 0.5.

##### 4.1.4.2 Scope

The benchmark deliberately fixes a number of choices to keep the number of confounded factors manageable. Trainable parameters are restricted to the six membrane parameters {*C*_*m*_, *g*_*L*_, *E*_*L*_, *V*_*T*_, Δ_*T*_, *V*_*r*_}. The adaptation parameters {*a, b, τ*_*w*_} are held at their ground-truth values during optimization; we were not able to fit them well (section 5.3) in any single-trace step-current experiments. In principle these parameters are observable through dedicated subthreshold ramps and periodic spike-evoking pulses [4], but those stimuli are absent from the protocol used here. By treating them as out-of-scope here, we can investigate what gradient-based fitting can do for membrane parameters.

Targets are synthetic: ground-truth and recovered parameters share the same forward model, so any failure to recover is attributable to the loss function and the optimizer rather than to model–data mismatch.

Stimuli are single step currents (5 ms delay, 90 ms duration, 100 ms total simulation, Δ*t* = 0.025 ms), matching the configuration used in the per-loss training notebooks so that the benchmark and the worked examples speak about the same regime. Multi-trace optimization, although described in section 4.1.3.3, is held out of this benchmark due to the reasons described above.

##### 4.1.4.3 Methods

We compare five methods (table S1), chosen so that the comparison isolates three questions: *which differentiable loss best supports gradient-based recovery, how a strong derivative-free baseline performs on the most informative of those losses*, and *whether the differentiable approximations introduced for the gradient methods cost the derivative-free baseline anything in turn*. The three gradient methods (`grad_-mse, grad_guarino, grad_vanrossum`) share the same Polyak optimizer configuration, surrogate kernel, and parameter reparameterization; they differ only in the loss they minimize. The fourth method (`nm_guarino`) runs Nelder–Mead on the soft Guarino objective, evaluating the same loss function as `grad_guarino` but ignoring its gradient. This pairing makes the gradient-vs-derivative-free comparison a like-for-like contrast on a single loss surface, rather than a comparison of two methods that also differ in their objective. The fifth method (`nm_guarino_hard`) runs Nelder–Mead on the *hard-feature* Guarino objective: the eight features are computed by exact threshold crossings on the simulated and target traces, with no temperature, sigmoid sharpness, or soft validity gates. This is the loss Guarino et al. [19] themselves use with an evolutionary-algorithm pipeline; including it lets us separate *the cost of soft-feature approximations to the derivative-free baseline* from *the cost of the surrogate gradient to the gradient methods*, which the soft `nm_guarino` alone cannot do.

##### 4.1.4.4 Paired directions, grid and sample sizes

Within a scenario, a single set of *N*_dir_ = 30 unit-sphere directions is drawn once from a fixed RNG seed and reused across all (*δ*, method) cells; the same direction *u*_*k*_ therefore generates the initialization *θ*_0_(*δ, k*) at every *δ*, and every method is evaluated on that identical initialization. Cross-method comparisons at fixed (scenario, *δ, k*) are therefore matched on the underlying perturbation: when one method succeeds and another fails on the same direction, this is a per-instance difference in optimization behaviour and no sampling artefact.

We use *δ* ∈ {0.02, 0.05, 0.1, 0.2, 0.4, 0.8}, spanning from near-recovery (~ 2% of bound width, no clipping) to perturbations large enough that most initializations are pinned to at least one parameter bound (~ 90% of runs at *δ* = 0.8; see table 1). The headline curves should be interpreted as basin-of-attraction estimates at small *δ* and as feasibility-projected recovery at large *δ*.

**Table 1.** Fraction of recovery-benchmark initializations with at least one parameter clipped to a bound, as a function of perturbation scale *δ*. Averages are over the 90 runs per *δ* (3 scenarios × 30 directions). At large *δ* most initializations lie on a face of the feasibility box rather than on the sphere of radius *δ* around *θ*^*\**^; the headline curves should be read as basin-of-attraction estimates at small *δ* and as feasibility-projected recovery at large *δ*.

| $\delta$ | % runs clipped | mean # clipped<br>(of 6 params) | max # clipped<br>(of 6 params) |
| --- | --- | --- | --- |
| 0.02 | 0.0% | 0.00 | 0 |
| 0.05 | 0.0% | 0.00 | 0 |
| 0.10 | 10.0% | 0.10 | 1 |
| 0.20 | 37.8% | 0.39 | 2 |
| 0.40 | 65.6% | 0.77 | 2 |
| 0.80 | 90.0% | 1.62 | 4 |

Three scenarios (`tonic, adaptation, initial_-bursting`) cover the qualitatively distinct firing patterns over which an AdEx fit must generalize. Total: 5 × 3 × 6 × 30 = 2700 runs per benchmark execution. For each (scenario, *δ*) tuple we compare Γ distributions between methods using the two-sample Kolmogorov–Smirnov test on the *N*_dir_ = 30 recovered Γ values per method.

#### 4.1.5 Subthreshold Recovery Benchmark

The recovery-radius benchmark of section 4.1.4 entangles two things that are hard to separate after the fact: the difficulty of the optimization problem, and the difficulty of getting any gradient at all through the discrete spike mechanism. A failure there could reflect either. To isolate the second factor we run a companion benchmark in the *non-spiking* regime, where the surrogate gradient never activates because no threshold crossing occurs. In the subthreshold regime, spike and adaptation parameters don’t contribute to the dynamics. Only three parameters shape the voltage trace — the membrane capacitance *C*_*m*_, the leak reversal *E*_*L*_ and the leak conductance *g*_*L*_. The task reduces to a three-parameter regression on a smooth voltage trace. Specifically, no custom-VJP (surrogate gradients) anywhere on the path from parameters to loss. This lets us ask a sharper question than the spiking benchmark can: with all spike-related problem aside, does gradient descent on the raw MSE loss actually behave worse than a derivative-free optimizer?

The construction reuses the perturbation machinery of section 4.1.4 with three changes. First, the stimulus is a subthreshold step (100 pA; 30 ms pre-stimulus baseline followed by a 100 ms step, Δ*t* = 0.025 ms), and a non-spiking guard rejects and resamples any perturbed ground truth whose target trace crosses threshold. In other words: we regenerate and disregard ground truths as long as they contain any spiking behaviour. This way every scenario stays in the subthreshold regime by construction. Second, all AdEx parameters are still perturbed to build a realistic unknown ground truth (section 4.1.4), but only {*C*_*m*_, *E*_*L*_, *g*_*L*_} are made trainable. Every fit starts from the same fixed initialization (the tonic subthreshold values with the spike parameters set to non-spiking defaults). Third, the two methods compared are gradient descent (Adam, learning rate 0.2, sigmoid parameter transform) and Nelder–Mead. Both minimize the identical MSE voltage loss, both are capped at the same budget of 25 loss evaluations: one Adam step against one Nelder– Mead function evaluation. A run counts as converged when a clean, non-surrogate re-simulation at the fitted parameters reaches an MSE of 0.1 mV^2^ or below.

We run one base parameter set (`tonic`) over the perturbation grid *δ* ∈ {0.1, 0.2, 0.3, 0.4} with 50 random directions each, giving 200 paired scenarios in which both optimizers fit the same target from the same initialization. Because the comparison is fully paired on the scenario, we report paired statistics: a Wilcoxon signed-rank test on each continuous metric (final loss, parameter-recovery error, evaluations-to-tolerance, wall-clock time) and a McNemar test on the binary converged/not outcome. Runs that never reach the tolerance within the shared budget are right-censored to the budget rather than discarded, so the speed comparison also credits the method that converges more often. The results are reported in section 5.3.3.

### 4.2 Compute environment

All benchmark and verification runs are executed on a single Apple MacBook Air (M1) with JAX configured to the CPU backend (`jax_platform_name=“cpu`”) for reproducibility across machines without an accelerator; gradient methods complete in ~1 s per run after Just-In-Time Compilation (JIT) warm-up, Nelder–Mead in ~5–10 s per run. The hard-feature variant `nm_guarino_hard` runs unjitted on every function evaluation (the spike-detection step uses host-side `numpy`), but a single forward simulation amortizes this so the per-run wall time stays in the same range as the soft NM variant. Total benchmark wall time is approximately 2 h. The hyperparameters in table S1 are the configurations validated in the per-loss training notebooks of section 5.3; no per-scenario tuning is applied within the benchmark.

### 4.3 Implementation

This section builds the tool the rest of the thesis relies on: a differentiable AdEx model for `Jaxley`. A simplified spiking neuron of this kind — with a discrete reset and exponential spike initiation — is not something `Jaxley`, otherwise designed for HH-type models, previously provided, so building it is the first of this work’s contributions. We construct it in two parts: the AdEx channel and its discrete-time formulation (section 4.3.1), and the surrogate-gradient variant — the soft reset and the choice of surrogate function — that makes the discrete spike mechanism differentiable (section 4.3.2). Both are verified against the Brian2 simulator and compared on runtime; the corresponding numerical results appear in section 5. The same construction pattern carries over to other simplified models such as LIF and Izhikevich, so the contribution is not limited to AdEx.

#### 4.3.1 AdEx Model Implementation

We implement the AdEx model as a `Jaxley` channel, following the formulation by Brette and Gerstner [4] (see eqs. (1) to (3)). Unlike HH-type channels that contribute currents to a shared voltage solver, simplified models like AdEx handle voltage integration directly within their `update_states()` method. We follow the implementation pattern of [5].

##### 4.3.1.1 Numerical Integration

The AdEx model consists of two coupled differential equations requiring different numerical treatment:

###### Adaptation current *w*

Equation (2) is a linear dynamic system in *w*, allowing the use of exponential Euler integration. It has the analytical solution

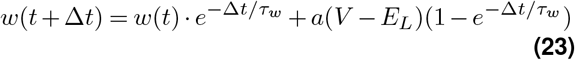

which gives increased numerical stability compared to e.g. forward Euler methods.

###### Membrane potential *V*

The exponential term in eq. (1) makes this equation nonlinear, requiring numerical integration, for example forward Euler:

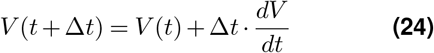

To prevent numerical overflow from the exponential spike mechanism, we clamp the argument:

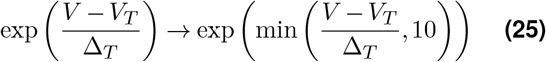

##### 4.3.1.2 Spike Detection and Reset

After each integration step, we check for threshold crossing (*V > v*_threshold_). If the threshold is crossed, we apply the reset conditions from eq. (3). For standard LIF, `AdEx` or `Izhikevich`, this uses `jax.lax.select(`) for conditional assignment. In the differentiable variants, we use the continuous formulation described in section 4.3.2.1 for both *V* and *w*:

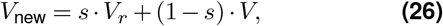

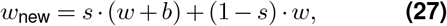

where *s* = *H*(*v* −*v*_th_) is the spike indicator (binary in the forward pass; its backward derivative is supplied by the surrogate *φ*_*β*_).

##### 4.3.1.3 Model Parameters

Table 2 summarizes the AdEx parameters and their default values. Some parameters have direct physiological counterparts (e.g., *C*_*m*_ or *E*_*L*_) and could technically be measured. However, to accurately reproduce neuron spiking behaviour, values can significantly diverge from biophysically plausible value ranges. This is expected and accepted when using simplified models, however, increases the demand of fitting strategies.

**Table 2.** AdEx model parameters and default values.

| Parameter | Symbol | Default | Unit | Trainable |
| --- | --- | --- | --- | --- |
| Membrane capacitance | $C_m$ | 200 | pF | Yes |
| Leak conductance | $g_L$ | 10 | nS | Yes |
| Leak reversal | $E_L$ | -70 | mV | Yes |
| Threshold potential | $V_T$ | -50 | mV | Yes |
| Slope factor | $\Delta_T$ | 2 | mV | Yes |
| Reset potential | $V_r$ | -58 | mV | Yes |
| Detection threshold | $v_{\text{threshold}}$ | 0 | mV | No |
| Adaptation time constant | $\tau_w$ | 30 | ms | Yes<sup>†</sup> |
| Subthreshold adaptation | $a$ | 2 | nS | Yes<sup>†</sup> |
| Spike-triggered adaptation | $b$ | 0 | pA | Yes<sup>†</sup> |
| Stimulation current | $I$ | 500 | pA | No |

More on trainability of the parameters in section 5.3.

***Point-Neuron Geometry Bridge*** The AdEx model is a point neuron defined with absolute units (*C*_*m*_ in pF, *g*_*L*_ in nS, currents in pA). Jaxley, however, is designed for geometry-aware biophysical simulations and operates with specific capacitance (*µ*F/cm^2^), current densities (*µ*A/cm^2^), and compartment surface areas. To bridge this mismatch, we assign a fixed fictitious cylindrical geometry with surface area *A* = 100 *µ*m^2^ (radius = 1 *µ*m, length = 50*/π µ*m) and store the absolute capacitance *C*_*m*_ (in pF) directly in Jaxley’s `capacitance` parameter. Why does this work? As stated in section 4.3.1, there are two different options on how voltage equations are solved in `Jaxley: update_states` and `compute_current`. Since `compute_current` is set to 0, all internal dynamics are handled entirely in `update_states`, which computes d*V/*d*t* = (−*I*_leak_ + *I*_exp_−*w*)*/C*_*m*_ in point-neuron units (pA / pF = mV/ms). This way, there is no double integration. However, external current stimulation is passed into the channel by `Jaxley’s convert_point_process_to_distributed`. *I*_nA_*/A* × 10^5^ [*µ*A/cm^2^]. With *A* = 100 *µ*m^2^ this simplifies to *I*_nA_ × 1000 = *I*_pA_ (numerically). The voltage solver then divides by capacitance: *I*_pA_ */ C*_*m*_ [mV/ms], which is in correct point-neuron units.

Importantly, *C*_*m*_ **is trainable**. Because the geometry is fixed, changing *C*_*m*_ during optimization only affects the two divisions described above — exactly the physical role of membrane capacitance. Therefore no geometry recalculations are needed. Conversion complexity is reduced to a simple unit change: *I*_nA_ = *I*_pA_ */* 1000, no parameter-dependent scaling is required.

##### 4.3.1.4 Implementation Verification

Before any of the gradient-based experiments make sense, we need to be confident that the forward simulation itself is correct. Otherwise, every later observation about loss landscapes, gradients, or training failures would be ambiguous: a real property of gradient-based optimization, or just a bug in the dynamics? To rule out the latter, we compare our AdEx channel against Brian2 [28], a widely-used and well-tested simulator for spiking neural networks. Both simulators are configured (almost^*^) identically:

- Time step: Δ*t* = 0.01 ms
- Simulation duration: 500 ms
- Stimulation duration: 400 ms
- Initial conditions: *V* (0) = *V*_*r*_, *w*(0) = 0

We evaluate three parameter configurations from Brette & Gerstner [4] and from Naud et al. [2] that produce distinct firing patterns:

###### Tonic spiking

Regular spiking without spike-frequency adaptation, serving as a baseline case. The parameters are the default parameters (see table 2).

###### Adaptation

Strong spike-triggered adaptation with slow decay, producing decreasing firing rates over time. The parameters used are *C*_*m*_ = 200, *g*_*L*_ = 12, *E*_*L*_ = −70, *v*_*T*_ = −50, Δ_*T*_ = 2, *v*_*r*_ = −58, *v*_threshold_ = 0, *τ*_*w*_ = 300, *a* = 2, *b* = 60, *I* = 500.

###### Original parameters

The original parameter set from Brette and Gerstner, demonstrating combined adaptation mechanisms. The parameters used are *C*_*m*_ = 281, *g*_*L*_ = 30, *E*_*L*_ = −70.6, *v*_*T*_ = −50.4, Δ_*T*_ = 2, *v*_*r*_ = −70.6, *v*_threshold_ = 20, *τ*_*w*_ = 144, *a* = 4, *b* = 80.5, *I* = 2500.

For each configuration we look at three things: an overlay of the voltage traces from both simulators as a qualitative sanity check, the number of spikes each simulator produces, and the maximum pair-wise difference in spike times,

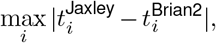

as a quantitative measure of agreement. The actual numbers are reported in section 5.1.

##### Runtime Performance Comparison to Brian2

We evaluate runtime performance of the AdEx implementation by comparing it to Brian2. To establish a fair comparison, we benchmark against the different available runtimes, namely the python environment using *cython* and c++. In the last case, the model gets compiled into native c++ code which then is compiled to the target platform. We perform all tests on an Apple MacBook Air (M1), see more details in section 4.2.

The benchmark is set up as follows: we simulate the AdEx equations with the same (tonic) parameters for different simulation lengths. Specifically, we time a single point neuron over a sweep of simulation durations (200–5000 ms, i.e. 8,000–200,000 integration steps) and separate the one-time compile/code-generation cost from the per-run integration walltime, which is the quantity compared. Each scenario is performed 20 times for a time step size of *dt* = 0.025. Brian2 is using the same numerical scheme for both equations, whereas Jaxley uses forward Euler for the voltage equation and exponential Euler for the adaptation.

This choice is made, to further speed up the Brian2 simulation. Results are presented in section 5.1.1.

#### 4.3.2 Integration of Surrogate Gradients into Jaxley

`Jaxley` is built on top of `JAX` and leverages its automatic differentiation capabilities for gradient-based parameter optimization. Neuron dynamics in Jaxley are defined through Channel classes, which specify three core methods: `update_states()` for state variable evolution, `compute_current()` for membrane current contributions, and `init_state()` for initialization of state variables. During simulation, JAX traces these functions to construct a computational graph, enabling automatic gradient computation and backpropagation. The discrete reset of eq. (3) breaks this graph: as discussed in section 2.3.2, the Heaviside derivative is zero almost everywhere, so gradients through spike events vanish.

##### 4.3.2.1 Implementation Details

To overcome this limitation, we employ surrogate gradients [16]: the forward pass uses the original Heaviside function to maintain correct spike dynamics, while the backward pass substitutes its derivative with a smooth surrogate. We implement this in `JAX via jax.custom_vjp`, which attaches a hand-written vector–Jacobian product to the otherwise non-differentiable spike indicator. All three surrogates introduced in section 2.3.3 — sigmoid (eq. (6)), exponential (eq. (7)), and SuperSpike (eq. (8)) — are available as drop-in choices; unless stated otherwise, experiments use the sigmoid surrogate with *β* = 5.0.

We further reformulate the conditional reset from a discrete selection

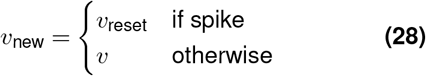

to a continuous, differentiable form

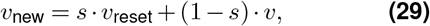

where *s* = *H*(*v*− *v*_th_) is the spike indicator. In the forward pass *s* is 0 or 1 exactly (the hard Heaviside *H*), so this expression is numerically equivalent to the discrete selection above and the recorded spike trace *s*(*t*) ∈ {0, 1} is binary — the continuous-blend form exists only so that JAX’s `custom_vjp` can route a smooth gradient (the surrogate derivative *φ*_*β*_) through the reset on the backward pass.

This approach can easily be applied to the existing LIF (`FireSurrogate`) and Izhikevich models in `Jaxley`, not only to the newly implemented AdEx model (`AdExSurrogate`).

##### 4.3.2.2 Implementation Verification

To verify the surrogate gradient implementation, we need to show what the surrogate is actually for: restoring non-zero gradients through spike events. We construct a minimal test case modelled after the demonstration in Neftci et al. [16] and split it into two checks. First, *forward equivalence*: by construction of `custom_vjp` the surrogate slope *β* enters only the backward rule, so `AdExSurrogate` and the standard `AdEx` channel evaluate the same hard Heaviside and same hard reset. We confirm this empirically by running both channels on the three Naud parameter sets and comparing spike counts and spike times.

Second, *gradient existence*. A short simulation is run with a step current strong enough to elicit several spikes, and we use the negative spike count ℒ = – ∑_t_*s(t)* as a toy loss; for each surrogate variant (sigmoid, exponential, SuperSpike) we compute *∂ℒ /∂θ* under `AdExSurrogate` and compare against the same gradient computed with the standard `AdEx` channel. The recorded spike output is the indicator 1[*v* ≥ *v*_th_], which is piecewise constant in *v*, so under standard autodiff the hard channel necessarily returns a structural zero with respect to *every* parameter — any non-zero gradient under the surrogate channel therefore demonstrates that the `custom_vjp` rule is wired correctly and that gradients propagate through spike events as intended. We pick `v_threshold` as the trainable parameter because its expected sign is unambiguous: *∂ ℒ/∂v*_th_ *>* 0, since raising the detection threshold drops spikes and increases the loss. Figure 2 shows the three surrogate kernels overlaid on the hard threshold; numerical results are reported in section 5.1.2.

Together, the two checks establish what the implementation is supposed to do: the forward pass behaves indistinguishably from the standard AdEx model, while the backward pass replaces a structurally zero gradient with a finite one. Whether the resulting gradients are *useful* for optimization — their magnitude, their support over the voltage range, and their behaviour under long simulation horizons — is a separate question, and one that the rest of this thesis is largely concerned with.

## 5 Results

This chapter reports four classes of empirical results, bearing on the thesis’s central question. Section 5.1 establishes that the forward model is correct, by comparing our `Jaxley` AdEx channel and its surrogate-gradient variant against Brian2 on three reference parameter sets. This is a precondition for the rest of the chapter. It rules out the possibility that any later failure is an artefact of the dynamics rather than a property of the optimization. We evaluate runtime performance of the AdEx implementation in `Jaxley` against Brian2 in section 5.1.1. Section 5.2 characterizes the loss surfaces the optimizer actually descends. Section 5.3 reports what happens when an optimizer is run on those surfaces: two worked examples (one synthetic, one on a real recording), a recovery-radius benchmark over 2,700 paired runs that compares three differentiable losses against two Nelder–Mead baselines on identical perturbed initializations and a companion subthreshold benchmark that isolates gradient descent from the spike mechanism entirely.

### 5.1 An AdEx Channel for Jaxley

The first concrete outcome of this thesis is a working AdEx channel for `Jaxley`, validated against an established reference simulator. Following the protocol of section 4.3.1.4, we compare our implementation to Brian2 [28] on all three reference parameter sets from Naud et al. [2] — tonic, adaptation and original — which together cover the qualitatively distinct firing regimes AdEx is meant to reproduce.

Across all three configurations, the two simulators produce identical spike counts, and the maximum pair-wise spike-timing difference stays below 1 ms. Figure 3 shows the side-by-side comparison of voltage traces, adaptation current *w*, and spike times. The remaining sub-millisecond differences are what one would expect given that the two simulators use different integration schemes for the membrane equation, and they are well below the temporal resolution at which we later evaluate spike-timing accuracy via the coincidence factor Γ.

**Figure 3.**
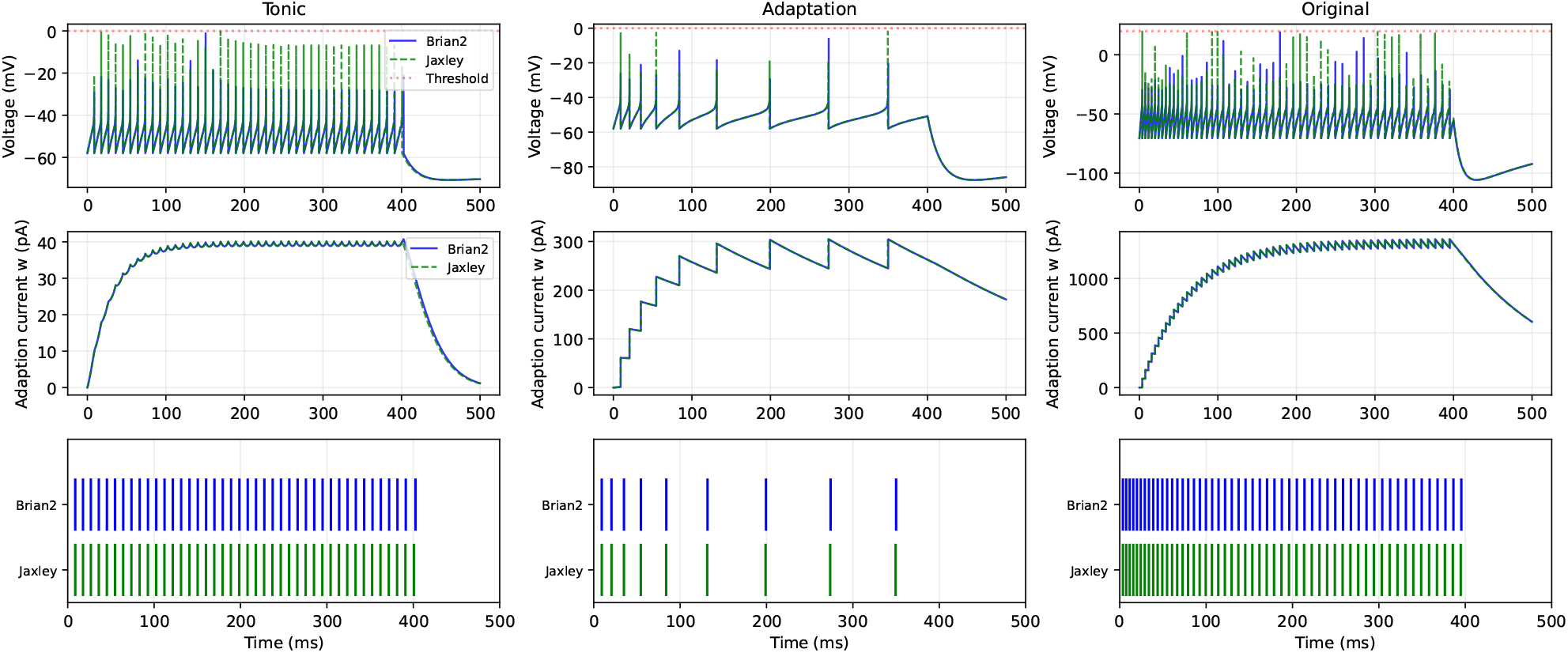
Comparison of our `Jaxley` AdEx implementation against Brian2 across three reference parameter sets. Left to right: tonic, adaptation and original parameters. Tonic and adaptation parameters refer to the parameters provided by [2], original parameters are the parameters used in the original AdEx paper [4]. Top row shows the simulated voltage traces; note that when the threshold is reached, the voltage is reset within the same time step — a behaviour that becomes important when thinking about loss functions later on. Second row displays the adaptation current *w*. Last row compares the spike timings between `Brian2` and `Jaxley`.

This validation matters for two reasons. First, it establishes that an AdEx-style simplified neuron — with its discrete reset and exponential spike initiation — can be expressed cleanly inside a framework otherwise designed for HH-type compartmental models. The same implementation pattern carries over to other simplified models such as LIF and Izhikevich, so the contribution is not strictly limited to AdEx. Second, and more importantly for the rest of this chapter: every optimization failure we report from here on can be discussed as a property of gradient-based training and loss-function design, rather than as a possible artefact of the underlying dynamics. The forward model is not the bottleneck.

#### 5.1.1 Runtime Comparison

Having established that the `Jaxley` AdEx channel reproduces Brian2’s dynamics, we ask what it costs to run. We follow the experimental setup of section 4.3.1.5. The spike counts logged alongside each timing (last column of table 3) confirm that the two simulators integrate the same model throughout: the largest disagreement anywhere in the sweep is 1 spike out of 93 (≤ 1.1 %) It grows with trace length, consistent with slow phase drift between two different and independent but individually converged integrators and is within expected error bounds and not attributed to model discrepancy.

**Table 3.** Per-run AdEx integration walltime (mean over 20 repeats, Δ*t* = 0.025 ms). Brian2 runs in its default runtime/Cython mode with its fastest scheme `(euler`, selected from a sweep over `euler, heun, milstein, rk2, rk4`); Jaxley uses forward Euler. The final column reports spike counts (Jaxley/Brian2) as an equal-model guard.

| $T$ (ms) | steps | Jaxley (ms) | Brian2 (ms) | speedup | spikes (Jx/Br) |
| --- | --- | --- | --- | --- | --- |
| 200 | 8000 | 0.462 | 142.53 | 308× | 10/10 |
| 500 | 20000 | 1.129 | 246.64 | 218× | 41/41 |
| 1000 | 40000 | 2.269 | 393.18 | 173× | 93/92 |
| 2000 | 80000 | 4.540 | 682.87 | 150× | 196/195 |
| 5000 | 200000 | 11.251 | 1489.74 | 132× | 507/505 |

In this default mode Jaxley is faster at every duration, by a geometric-mean factor of ×188 (range 132–308×; table 3). The two simulators scale linearly in the number of integration steps, as expected for a single uncoupled compartment. A per-step cost fit (fig. 4) gives 0.056 *µ*s/step for Jaxley against 6.96 *µ*s/step for Brian2 — a 124 × gap per timestep — with Jaxley’s log–log scaling exponent at 0.99 (*R*^2^ = 1.00). Brian2’s apparent exponent of 0.73 is not sublinear integration but the signature of a fixed per-run dispatch overhead of tens of milliseconds that dominates the shorter traces. Expressed as throughput, Jaxley sustains ≈440× real time across the whole sweep, whereas Brian2 in this mode reaches at most 3.4× real time on the longest trace. The one-time cost does not change the picture: Jaxley’s JIT compile (0.13 s mean) is smaller than Brian2’s code generation (0.70 s mean), so Jaxley wins even for a single run including overhead.

**Figure 4.**
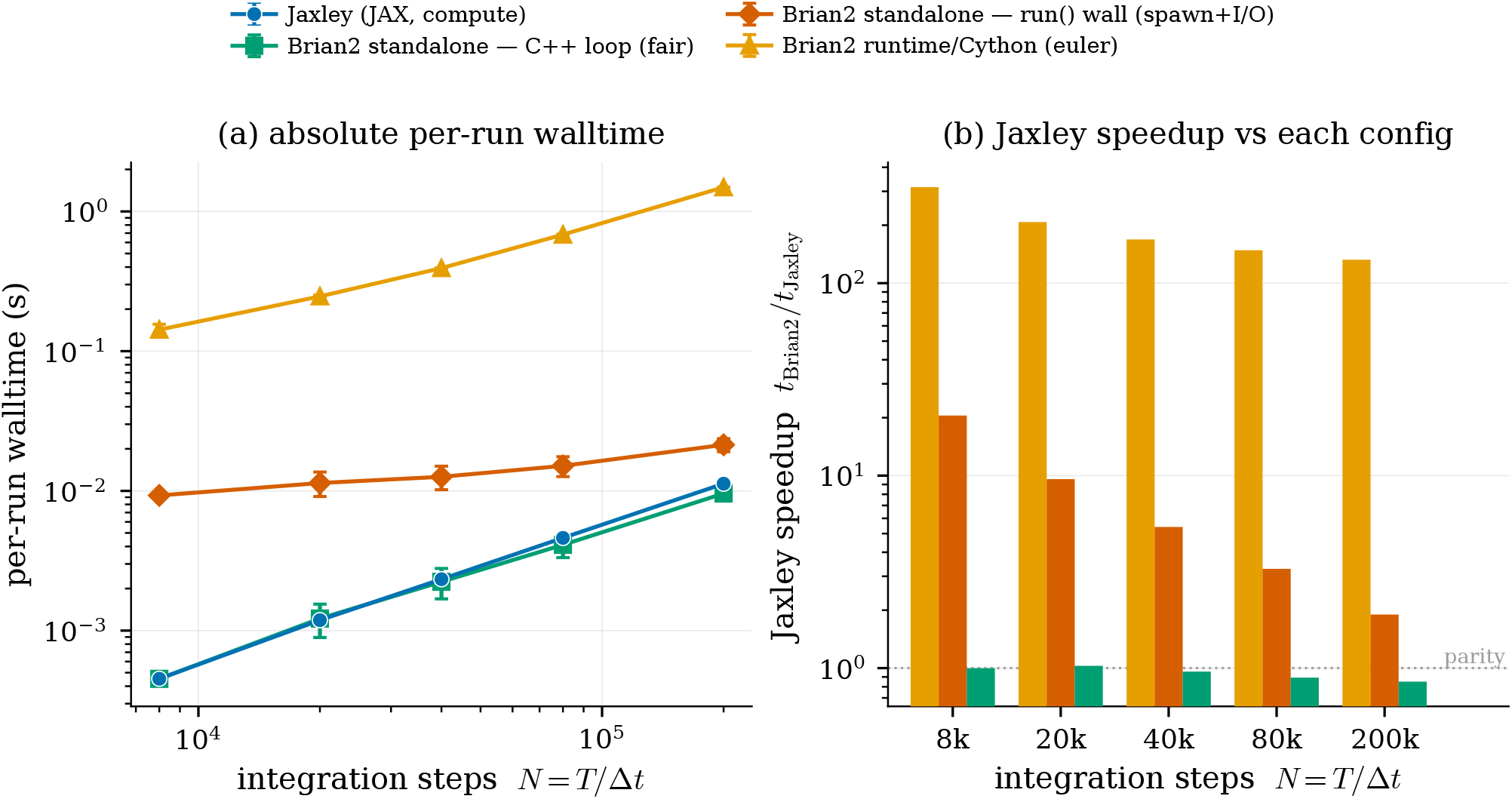
Per-run AdEx integration walltime across the full spectrum of Brian2 configurations, as a function of the number of integration steps *N* = *T/*Δ*t* (error bars are ±std over 20 repeats, often smaller than the markers). **(a)** Absolute walltime (log–log): Jaxley (forward Euler, warm in-process compute) against Brian2 in its default runtime/Cython mode and its best-case compiled `cpp_standalone` device. *C++ loop* is the standalone binary’s self-reported integration time; *run() wall* additionally pays per-call process spawn and result I/O. All curves scale linearly in *N*; the vertical offset to the runtime/Cython curve is the ~ 10^2^× per-run gap of table 3. **(b)** The corresponding Jaxley speedup *t*_Brian2_*/t*_Jaxley_ (log *y*) against each configuration.

The ~ 10^2^ × headline should not be over-read, and fig. 4(a) is included to keep the comparison honest. Brian2’s default runtime mode keeps its integration loop in the Python interpreter, which for a single point neuron is dominated by per-timestep interpreter overhead — Brian2’s worst case. Its `cpp_standalone` device compiles the whole simulation to a standalone C++ program with no Python in the loop, using Brian2’s own default optimized build flags. Against that best case the fair compute-vs-compute comparison is essentially a tie: Jaxley ranges from 1.0× at short durations to 0.9× (i.e. marginally slower than the C++ loop) at 5000 ms. The end-to-end `run()` wall of the standalone binary, which re-executes the compiled program and pays OS process spawn plus result-file I/O on every call, sits between these extremes at 1.9– 20.5 ×. The defensible statement is therefore that the Jaxley AdEx implementation is between roughly on par with (compiled C++) and two orders of magnitude faster than (default runtime) Brian2, depending entirely on Brian2’s configuration. The point of this benchmark is not that Jaxley out-computes an optimized C compiler on a scalar loop — it does not — but that it reaches comparable raw throughput while remaining end-to-end differentiable, batchable via `vmap`, and GPU-portable.

#### 5.1.2 Surrogate Channel Verification

We run the two-part verification protocol of section 4.3.2.2. On all three Naud parameter sets, the forward pass of `AdExSurrogate` matches the standard `AdEx` channel perfectly. Both voltage trace and spike times are identical as can be seen in fig. S5. For the gradient-existence check, both channels report ℒ = − 21 on the tonic set at the operating point; the gradients are summarized in table 4.

**Table 4.**
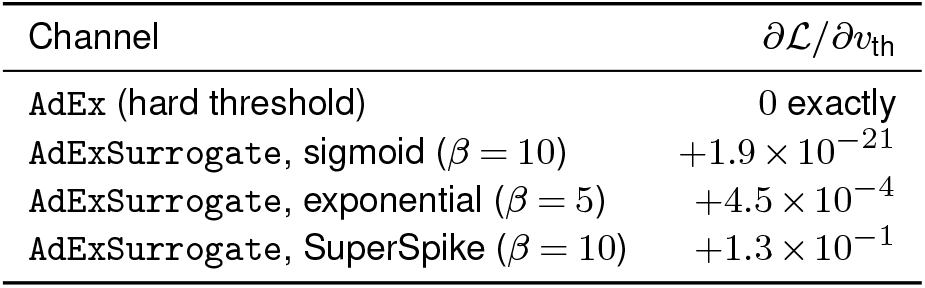
Gradient of the negative spike count with respect to v_threshold on the tonic parameter set (ℒ = −21 in all rows).

| Channel | $\partial \mathcal{L} / \partial v_{\text{th}}$ |
| --- | --- |
| AdEx (hard threshold) | 0 exactly |
| AdExSurrogate, sigmoid ($\beta = 10$) | $+1.9 \times 10^{-21}$ |
| AdExSurrogate, exponential ($\beta = 5$) | $+4.5 \times 10^{-4}$ |
| AdExSurrogate, SuperSpike ($\beta = 10$) | $+1.3 \times 10^{-1}$ |

The hard channel returns a structural zero; all three surrogates return a finite gradient of the correct sign. At a fixed operating point the magnitudes span more than twenty orders of magnitude — a property of the surrogate family alone (cf. fig. 2) — which couples directly to learning-rate sensitivity in section 5.3.

### 5.2 Loss Function Analysis

Before we report parameter-recovery numbers in section 5.3, this section characterizes the geometry of the loss surfaces. The motivation is diagnostic: a flat success/failure number does not say *why* the method fails. The recovery-radius benchmark in section 5.3.2 will show that all three differentiable losses fail — they do so for different reasons. We therefore look at the shape of each loss directly: the position of the spiking valley, the extent of the non-spiking plateau, and the direction of the gradient on that plateau. The geometric features predict the failure modes the benchmark later confirms.

For every pair of trainable parameters we evaluate four differentiable losses (MSE, SDTW, the Guarino feature-based loss, and the Van Rossum distance) on a 20 × 20 grid spanning the parameter bounds. The remaining parameters are held at the Naud `adaptation` reference values of section 4.3.1.4. The same grid evaluation produces a paired gradient field *∇*_*θ*_*ℒ* via JAX autodiff. Gradient computations were performed using a sigmoid surrogate with *β* = 5.0. The magnitude is displayed for five different parameters (for compute time reasons). Loss and gradient panels can be found in section A.2;

#### 5.2.1 Shared structure across losses

Three features recur across every loss surface we plot (section A.2):

- **Narrow low-loss valleys**. The spiking region forms thin, curved valleys rather than isotropic (round, bowl shaped) basins. Valleys are often parallel to one axis (e.g. *g*_*L*_–*E*_*L*_ trades off along a single direction), meaning small updates along the wrong axis leave the loss unchanged.
- **Large non-spiking plateaus**. Substantial regions of parameter space yield a constant (or near-constant) loss because no spikes are produced and the loss reduces to a target-only constant. Soft spike-detection terms supply a residual gradient on these plateaus, but the gradient is small and its direction is not constrained by the true distance to the spiking valley.
- **Plateau gradient magnitude, not direction**. On the spike-aware losses (Guarino, Van Rossum) the surrogate can supply a small but non-zero gradient on the non-spiking plateau via *φ*_*β*_(*V* −*V*_th_). The resulting gradient direction does not point necessarily towards the spiking transition, see fig. 6. In any case, its magnitude is several orders below the gradient inside the spiking region.

**Figure 5.**
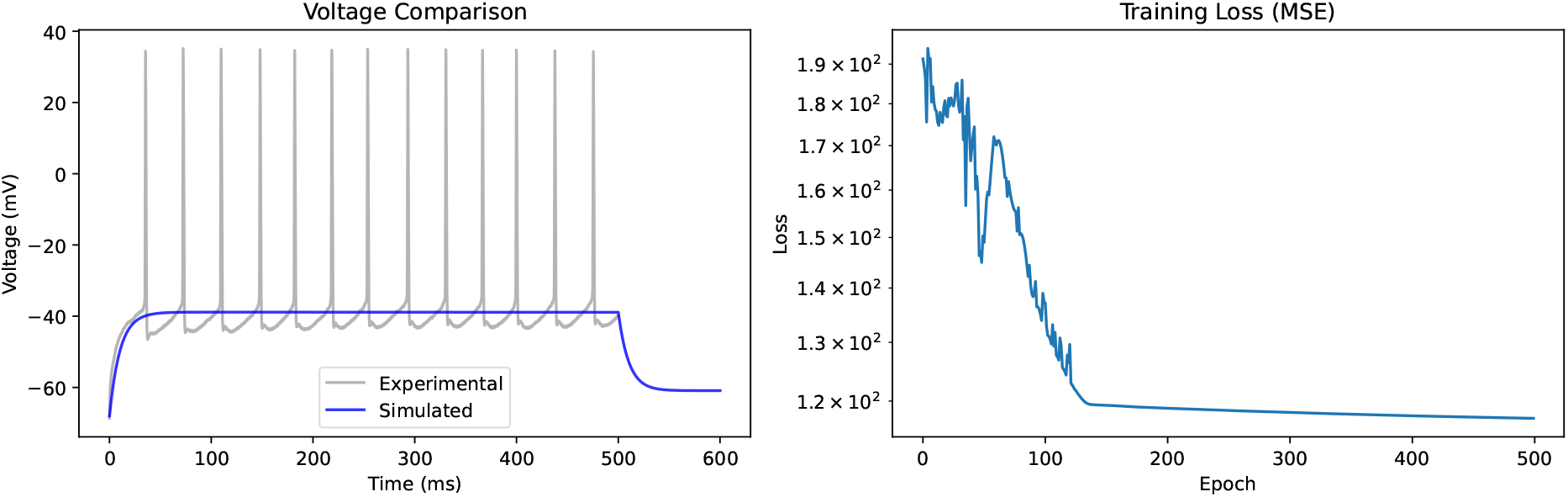
MSE training result on voltage recordings of a mouse striatal projection neuron. (a) The optimizer converges to a non-spiking subthreshold solution (blue) that tracks the experimental resting potential but produces no spikes. (b) The loss decreases monotonically, confirming that the non-spiking solution is a genuine local minimum of the MSE objective — eliminating spikes reduces point-wise error more effectively than aligning them.

**Figure 6.**
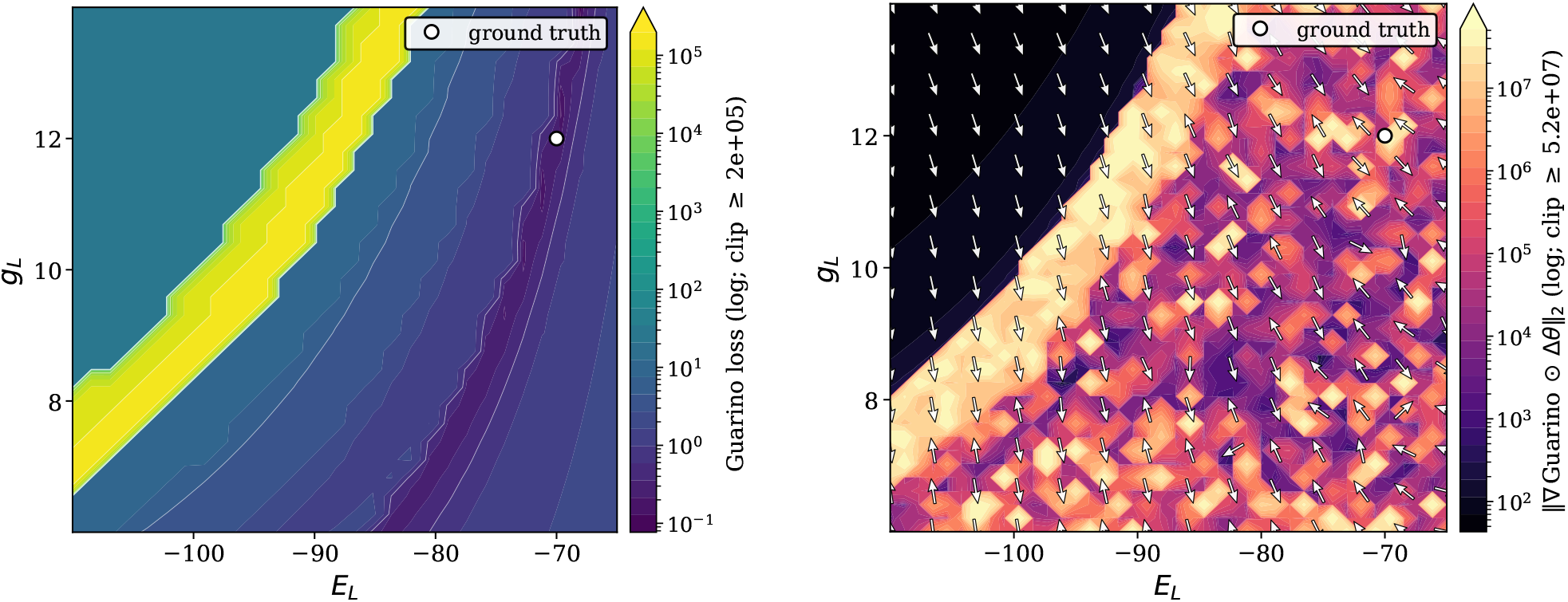
Guarino loss landscape (left) and bound-normalized gradient field (right) on the *E*_*L*_–*g*_*L*_ plane around the Naud `adaptation` reference point. SuperSpike surrogate, *β* = 10. Left: L_Guarino_ on a log scale spanning six orders of magnitude. Right: *∥∇*L ⊙ Δ*θ∥*_2_ on a log scale. Arrows show the unit direction of steepest *ascent* in bound-normalized coordinates; the optimizer steps opposite. The ground-truth marker sits inside the narrow curved valley of low loss (dark purple, left), bounded on the upper-left by the low spike frequency, high loss penalty ridge (yellow), The top left corner shows the plateau where the simulated trace loses all spikes. On the right panel the penalty ridge dominates the gradient magnitude (~ 10^7^). The plateau gradient outside the ridge is non-zero (~ 10^2^–10^3^ in bound-normalized units) and uniformly directional, pointing south-east towards the ridge.

The three subsections that follow show what these shared features look like in each loss individually. Before that a word about how gradient magnitudes vary between parameters.

##### Gradient magnitude asymmetry between parameters

There are huge differences between how susceptible to change different parameters are. Figure S6 shows the bound-normalized gradient field on the (*a, τ*_*w*_) plane around the Naud `adaptation` reference point (*a* = 2 nS, *τ*_*w*_ = 300 ms), under Van Rossum with SuperSpike *β* = 10. The quiver is dominated by the horizontal (*a*) component at essentially every grid cell, even after bound-normalization puts the two axes on the same dimensionless scale. This is a real geometric property, not the unit-scaling artefact. It is mostly explained by a structural asymmetry in how AdEx parameters enter the dynamics. One class (*E*_*L*_, *g*_*L*_, *C*_*m*_, *V*_*T*_, *a*) enters the right-hand side of the membrane or adaptation equation at every timestep, so a small perturbation rescales or shifts *V* (*t*) (or *w*(*t*)) at every point along the trajectory and accumulates loss sensitivity over the full integration window. A second class (*b, τ*_*w*_, Δ_*T*_) only feeds into the dynamics through narrow channels. *b* acts as a discrete kick on *w* at spike times, so its gradient is gated by spike events and vanishes whenever the cell is silent or near-silent. *τ*_*w*_ is a relaxation timescale whose effect is only visible in longer simulations. Δ_*T*_ enters an exponential whose value is dominated by *V*_*T*_ at most operating points. The bound-normalized gradient field inherits this asymmetry directly. Rescaled units cannot make *b* act outside spike events, nor make *τ*_*w*_ act on timescales the stim window does not visit. The (*a, τ*_*w*_) slice above is one instance of this pattern; the same imbalance recurs whenever a continuously-acting parameter is paired with an event-driven or timescale one, and it is one of the reasons *b* and *τ*_*w*_ are systematically harder to recover.

#### 5.2.2 Mean Squared Error: the non-spiking attractor

The MSE surface has visibly the largest number of local minima. Across all 28 parameter pairs the surfaces display thin, axis-aligned spiking valleys. We observe large magnitudes in parameter areas where Van Rossum and Guarino show small loss values. This indicates that error is aggregated based on small timing mis-alignments, meaning that small temporal mis-alignments with correct spike count produce high errors using the MSE loss and small errors using feature based losses.

This geometry has an immediate empirical signature in training: starting from a tonic spiking initialization, the MSE optimizer deletes the simulated spikes rather than aligning them, because deleting spikes is the easier to reach loss target. Figure 5 shows the resulting trajectory on voltage recordings of a mouse striatal projection neuron: the trace tracks the resting potential with no spikes, and the loss decreases monotonically across all 50 epochs. MSE’s plateau gradient therefore points *away* from the spiking valley rather than towards it.

#### 5.2.3 Guarino: feature artefacts everywhere

The Guarino feature-based loss replaces MSE’s point-wise voltage comparison with a weighted sum of relative errors on eight discrete spike features (section 4.1.2.2). This can be supplemented by a soft validity penalty whenever the simulated trace does not produce enough spikes to evaluate a feature. Both changes target the failure mode diagnosed for MSE: removing spikes is no longer cheaper than aligning them, because the missing-feature penalty contributes a fixed positive term to the loss whenever a feature cannot be evaluated. The geometry near a spiking ground truth reflects this redesign.

Figure 6 shows the loss and the gradient field on the *E*_*L*_–*g*_*L*_ plane around the Naud `adaptation` reference point, on a logarithmic scale that spans six orders of magnitude. A narrow curved valley (dark purple, ℒ ≲ 10^0^) contains the ground truth and traces the locus where the leak equilibrium and conductance jointly produce the correct spike count and timing. In the top left corner, the missing-feature penalty turns on as the simulated trace loses spikes; The plateau is bounded on the upper-left by a sharp yellow ridge (ℒ ~ 10^5^). This is an interesting artefact of the feature based loss we constructed: the non-spiking area attributes a fixed penalty per feature. Parameters within the yellow ridge area produce voltage traces with a single or very few spikes. This disables the fixed added penalty, and instead large errors (e.g. in spike frequency) lead to huge loss values — gradient signals on the plateau therefore *discourage* moving into the optimal direction. All of this behaviour is basically irrelevant for sampling-based losses, which indicates why Guarino performs well in the benchmark when combined with Nelder– Mead.

Overall we can make three observations from the right panel of fig. 6. First, *∥∇*ℒ ⊙Δ *θ∥*_2_ spans five orders of magnitude across this single slice: ~ 10^2^ in the plateau interior to ~ 10^7^ on the penalty ridge. Second, on the non-spiking plateau the gradient field is approximately uniform — arrows point in essentially the same south-east direction across a region tens of millivolts wide, and that direction is the descent direction towards the ridge, whereas on the other side of the ridge, gradient behaviour is a lot more chaotic, varying over multiple orders of magnitude between evaluated points. Third, the largest gradients sit on the penalty ridge itself, not at the ground truth. All of this highlights the difficulties when applying feature based losses to gradient methods. The sheer number of hyperparameter needed to successfully finetune this loss may simply be too high (features specific weights, penalties, smoothness parameters, …).

#### 5.2.4 Van Rossum: a smoother loss, still chaotic gradient

The Van Rossum distance (section 4.1.2.3) is the only loss in our comparison that supplies a smooth path from the spike train *s*(*t*) to the loss value: the exponential kernel *K* and the *L*^2^ norm are themselves differentiable, so no softness hyperparameter is required to make the loss path smooth. The geometric prediction is that the Van Rossum surface should be smoother in the spiking regime than Guarino’s — the kernel convolution averages over a *τ*_kernel_-wide window, so small spike-timing perturbations no longer produce discrete penalty-ridge transitions. The empirical picture in the bound-normalized gradient field is more nuanced.

Figure 7 shows the Van Rossum loss and gradient field on the same *E*_*L*_–*g*_*L*_ slice and surrogate configuration as the Guarino panel of fig. 6 (SuperSpike, *β* = 10), so the two figures differ only in the loss.

**Figure 7.**
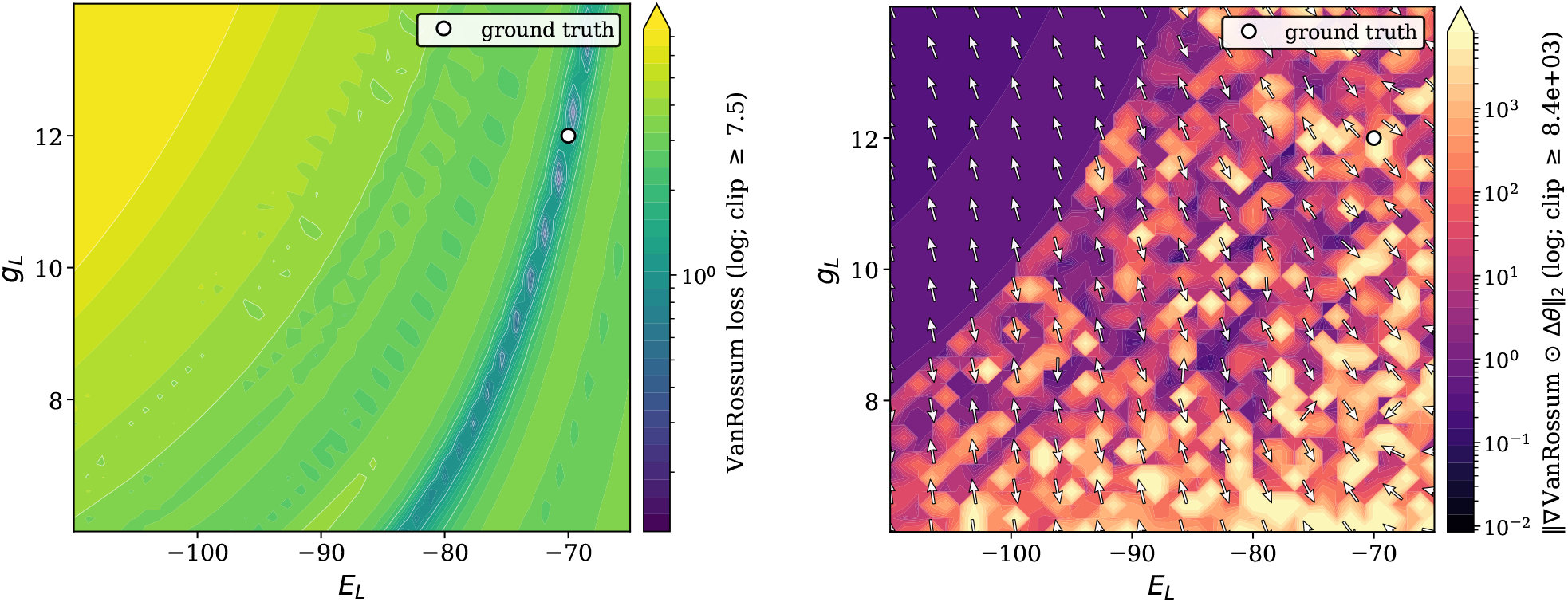
*E*_*L*_–*g*_*L*_ plane around the Naud `adaptation` reference point. Left: Van Rossum loss, Right: bound-normalized gradient-norm landscape with quiver overlay. SuperSpike surrogate, *β* = 10, matched to fig. 6. Left: 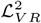 on a log scale spanning less than two orders of magnitude. Right: *∥∇*L⊙Δ*θ∥*_2_ on a log scale. Arrows show the unit direction of steepest *ascent* in bound-normalized coordinates; the optimizer steps opposite. The upper-left non-spiking plateau carries a small (~ 10^*−*1^) but directionally consistent ascent gradient. The lower-right spiking region exhibits a granular pattern of order-of-magnitude jumps between neighbouring grid cells: the surrogate’s narrow active window plus the binary *s*(*t*) mean that a single grid step can shift a spike in or out of the trace, producing a discrete change in *K* * *s*(*t*) and a step change in *∥∇*ℒ*∥*.

Two contrasts with the Guarino panel stand out. First, the feature penalty ridge of fig. 6 is absent: Van Rossum has no discrete penalty for losing a ‘feature’, so the loss climbs monotone and smoothly from valley to plateau across the diagonal transition rather than jumping by four orders of magnitude. This is aided by the kernel-smoothing effect the design argument predicts (section 4.1.2.3). Second, the bound-normalized plateau gradient is much smaller for Van Rossum (~10^−1^) than for Guarino (~10^2^): without the validity-flag penalty’s soft-gated contribution, the only signal on the plateau comes from *φ*_*β*_(*V* −*V*_th_) in the brief window when the leak dynamics carry *V* near threshold, which on the non-spiking plateau happens rarely or not at all. The kernel-smoothing benefit on the spiking branch is therefore paid for by a weaker plateau gradient on the non-spiking branch. This could be described as a magnitude bottleneck, and explains the non-moving parameters in non-spiking starting configurations.

The spiking region (lower-right of the diagonal) is *not* the smooth basin the kernel-convolution argument would suggest in isolation. Instead, it is dominated by a high-contrast checkerboard of bright and dark patches whose magnitudes differ by an order of magnitude between neighbouring grid cells. This might be explained by the binary indicator *s*(*t*): small parameter perturbation can shift one spike’s threshold-crossing across a single timestep and add or remove an event from the trace. The Van Rossum loss reads this as a discrete jump in *K* * *s*(*t*), and the autodiff backward pass attributes the jump via `custom_vjp` to the surrogate kernel at that timestep, producing a gradient at the affected grid cell that is large but uncorrelated with its neighbours’ gradients. The kernel convolution smooths the loss over timing perturbations of *existing* spikes; it does not smooth over spike-count changes, which is what dominates the region near and beyond the ground truth at any finite *β*.

##### Surrogate-family dependence

Figure 8 reports the same Van Rossum gradient field on the same slice for SuperSpike and sigmoid surrogates at three slopes *β* ∈ {5, 10, 25}. All six panels use identical axes, the same loss, and the same forward simulation; only the backward-pass surrogate kernel changes.

**Figure 8.**
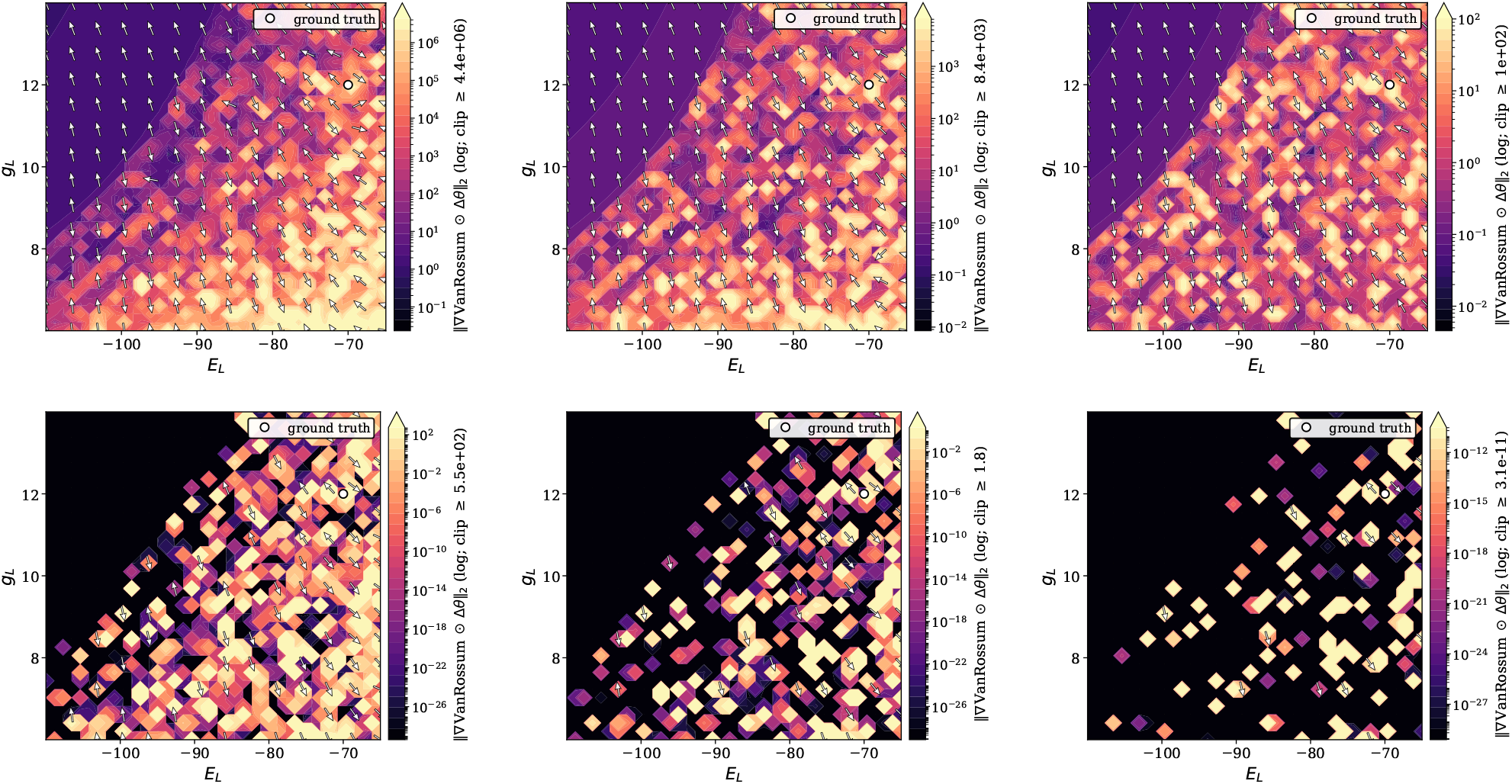
Van Rossum bound-normalized gradient field on the *E*_*L*_–*g*_*L*_ plane around the Naud `adaptation` reference point, swept over surrogate family and slope. Top row: SuperSpike (*β* = 5, 10, 25). Bottom row: sigmoid (*β* = 5, 10, 25). SuperSpike’s heavy tail 1*/*(*β* |*x*| + 1)^2^ produces gradient signal across the entire plane at every *β*: spiking-region magnitudes shrink from ~10^6^ at *β* = 5 to ~ 10^2^ at *β* = 25, but the spatial coverage is preserved. The sigmoid’s exponentially-suppressed *φ*_*β*_ (*x*) = *β σ*(*βx*)(1 − *σ*(*βx*)) collapses to numerical noise off-threshold: the dead zone (black) covers the non-spiking plateau at *β* = 5, most of the slice at *β* = 10, and the entire plane at *β* = 25 (magnitudes reach 10^*−*27^, well below denormalized float32 precision).

### 5.3 Parameter Recovery

We now turn from the geometry of the loss surfaces to what happens when an optimizer actually descends them. Throughout this section we follow the benchmark setup from section 4.1.4. The section is organized in two parts. Section 5.3.1 presents a single worked example in which gradient-based recovery succeeds somewhat; enough to establish that the approach is viable in principle, but already revealing that the problem is under-constrained. Section 5.3.2 then reports the recovery-radius benchmark: three differentiable losses against two Nelder–Mead baselines on matched initializations. The benchmark is the headline result of the chapter, and the picture it produces is uncomfortable: gradient recovery clears the no-optimization floor only at the smallest perturbations, and never matches Nelder–Mead on the same objective. The geometric features of section 5.2 suggest this pattern in advance.

#### 5.3.1 A Worked Example: Recovering a Tonic AdEx Cell

Before turning to the per-loss breakdown and the systematic benchmark, we present one configuration in which gradient-based fitting succeeds cleanly. The ground truth is the Naud tonic parameter set (section 4.3.1.4); the six trainable membrane parameters {*C*_*m*_, *g*_*L*_, *V*_*r*_, *V*_*T*_, *E*_*L*_, Δ_*T*_} are perturbed by approximately 15 % of their bound range, and the optimizer is run for 50 epochs of single-stage training under the Guarino feature-based loss (section 4.1.2.2). The adaptation parameters {*a, b, τ*_*w*_} are held at ground truth, matching the scope fixed for the recovery-radius benchmark in section 4.1.4.2. Figure 9 reports the outcome.

**Figure 9.**
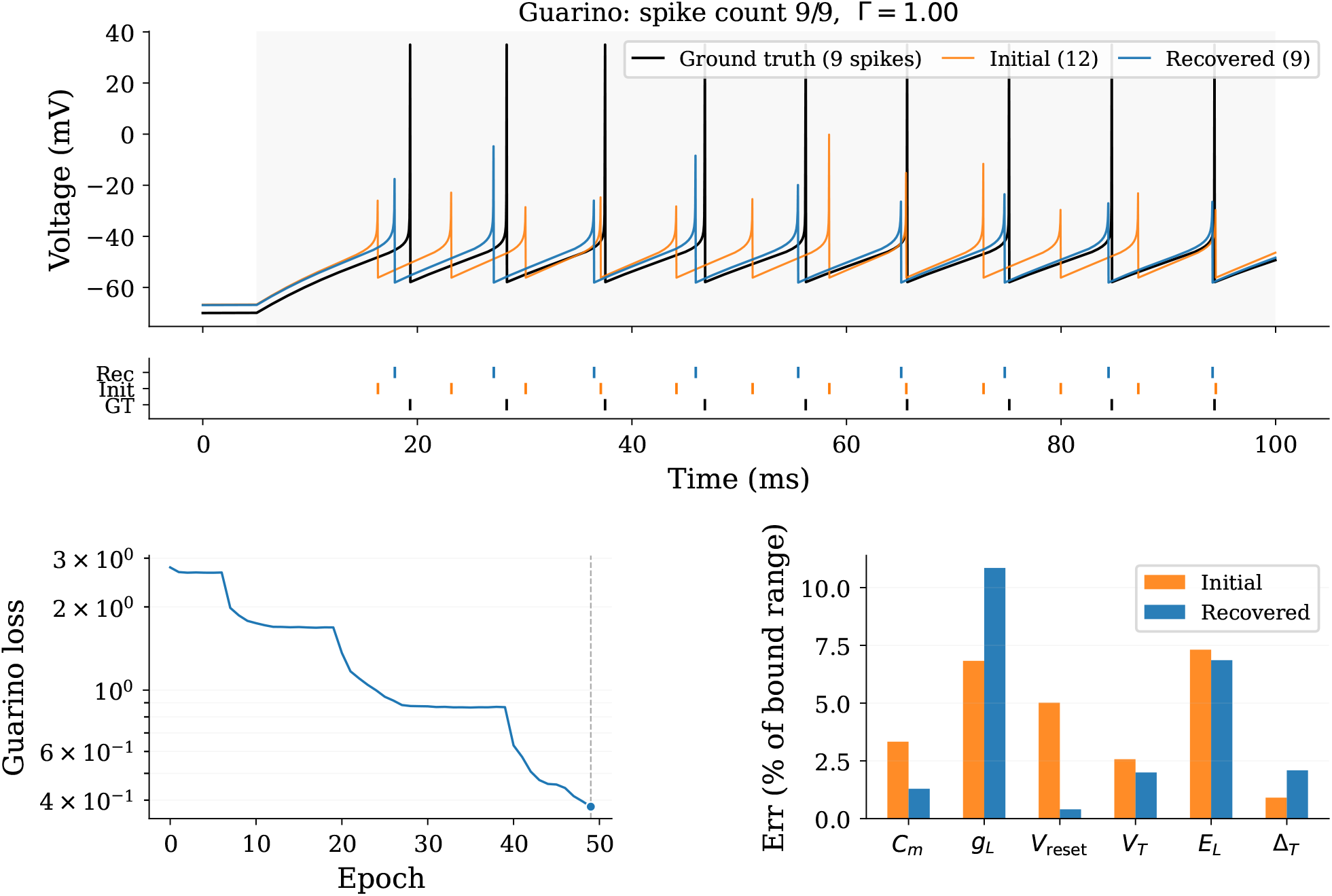
Single-stage recovery of a tonic AdEx cell under the Guarino loss. Top: voltage traces of the ground truth (black, 9 spikes), the perturbed initialization (orange, 12 spikes) and the recovered fit (blue, 9 spikes), with the corresponding spike raster below. Bottom left: training loss across 50 epochs; the dashed line marks the epoch at which the best parameters were retained. Bottom right: per-parameter absolute error before and after training, normalized by the width of the parameter bounds.

At the trace level the recovery is visually convincing. The perturbed initialization produces 12 spikes against the target’s 9 and misses most spike times by several milliseconds; after training the recovered trace matches the ground truth spike-for-spike and yields a coincidence factor of Γ = 1.00 at the Δ = 2 ms tolerance fixed in section 4.1.1. The loss curve descends monotonically over roughly one order of magnitude, with the steepest drop concentrated between epochs 35 and 45 where the optimizer successfully finds the right spike count.

The per-parameter panel tells a more nuanced story. Two of the six trainable parameters — *C*_*m*_ and *V*_r_ — tighten substantially towards their ground-truth values. However, other parameters change marginally or drift further away from the ground truth. Especially *g*_*L*_’s final error greatly exceeds the initial perturbation. The optimizer is therefore not recovering the parameter vector pointwise; it is settling into a local minimum combination that reproduces the target spike train while trading error across correlated axes. The loss-landscape evidence behind this compensation is taken up in section 5.2, and its implications for parameter identifiability are discussed in section 6.

This success is not generic. The same loss, the same hyperparameters and the same ground-truth parameter set fail to recover a comparable fit once the perturbation is increased, the firing regime is changed, or the adaptation parameters are made trainable. The recovery-radius benchmark of section 5.3.2 quantifies how quickly the success region closes; the geometric counterpart in section 5.2 explains why.

#### 5.3.2 Recovery-Radius Benchmark

The worked example of section 5.3.1 shows that gradient-based recovery can be possible; however, it does neither establish how robust the recovery is, nor how it compares against a derivative-free baseline on the same objective. The recovery-radius benchmark of section 4.1.4 answers both questions: it aggregates parameter recovery over 5 methods × 3 scenarios × 6 perturbations × 30 directions = 2700 paired runs, producing a recovery-success curve per method and a single recovery radius per (scenario, method) pair.

##### 5.3.2.1 Headline result: success vs. perturbation

Figure 10 shows *P* (Γ *>* 0.5) as a function of the perturbation scale *δ*, with one panel per scenario and one curve per method. A no-optimization floor (`init_floor`, the success rate of the perturbed initialization evaluated without any update) is included as a reference; methods below this floor have regressed.

**Figure 10.**
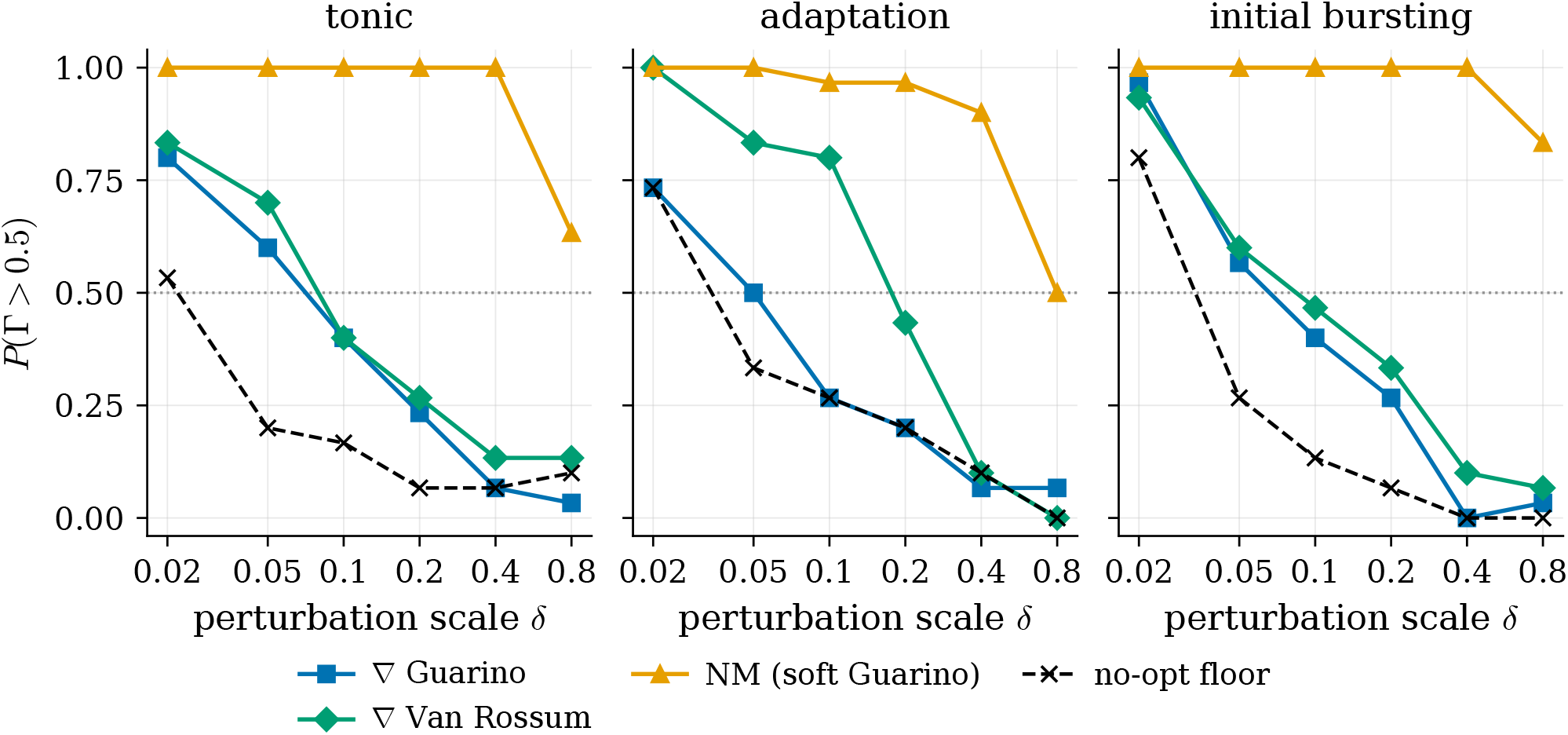
Recovery success *P* (Γ *>* 0.5) as a function of the relative perturbation scale *δ*, per scenario. Each curve aggregates 30 paired random-direction perturbations; methods are evaluated on identical (*θ*^*\**^ + *δu*_*k*_) initializations within each scenario. The Γ *>* 0.5 threshold follows section 4.1.1; the dashed horizontal at 0.5 marks the success-probability level used to define the recovery radius. `init_floor` reports Γ at the perturbed initialization with no optimization applied.

Four observations stand out. First, every gradient loss but MSE clears the floor at small *δ* (median Γ_0.05_ = 1.00 for `grad_vanrossum`), then collapses by *δ* = 0.1. Second, both Nelder–Mead variants dominate every gradient method on every scenario at every *δ*: they have the largest recovery radii and the highest success rates inside them. Third, `grad_mse` performed consistently the worst, consistent with the MSE landscape geometry of section 5.2.2. Fourth, the two Nelder–Mead variants are indistinguishable: `nm_guarino` (soft features) and `nm_guarino_hard` (exact features) report identical *δ*_50_ = 0.8 and median Γ_0.05_ = +1.00 on every scenario, with per-direction outcomes differing by at most one direction out of 30.

##### 5.3.2.2 Recovery radii

Table 5 reports two summary statistics per (scenario, method) pair: the recovery radius *δ*_50_ (the largest grid value of *δ* for which *P* (Γ *>* 0.5) ≥ 0.5) and the median Γ over all directions at *δ* = 0.05. The median at *δ* = 0.05 is the most diagnostic single number: *δ* = 0.02 is too easy (every reasonable method scores well) and *δ* ≥0.2 is too hard for the gradient methods (most rows are non-spiking failures dominated by the surrogate’s plateau gradient).

**Table 5.** Recovery-radius benchmark summary. *δ*_50_ is the largest grid value of *δ* ∈ {0.02, 0.05, 0.1, 0.2, 0.4, 0.8} for which *P* (Γ *>* 0.5) ≥ 0.5 over the 30 paired directions (*δ*_50_ < 0.02 written as “−” when even the smallest perturbation does not reach 50% success). Γ_0.05_ is the median Γ over directions at *δ* = 0.05. State-of-the-art Γ on real recordings is ≈0.82 [25]; the benchmark uses synthetic targets, so the achievable ceiling is Γ = 1.

| Method | tonic |  | adaptation |  | initial_bursting |  |
| --- | --- | --- | --- | --- | --- | --- |
| | $\delta_{50}$ | $\Gamma_{0.05}$ | $\delta_{50}$ | $\Gamma_{0.05}$ | $\delta_{50}$ | $\Gamma_{0.05}$ |
| `grad_mse` | – | 0.00 | – | 0.00 | – | 0.00 |
| `grad_guarino` | 0.05 | 1.00 | 0.05 | 0.54 | 0.05 | 0.90 |
| `grad_vanrossum` | 0.05 | 1.00 | 0.1 | 1.00 | 0.05 | 1.00 |
| `nm_guarino` | 0.8 | 1.00 | 0.8 | 1.00 | 0.8 | 1.00 |
| `nm_guarino_hard` | 0.8 | 1.00 | 0.8 | 1.00 | 0.8 | 1.00 |
| `init_floor` | 0.02 | 0.07 | 0.02 | 0.32 | 0.02 | 0.01 |

##### 5.3.2.3 What the benchmark does not measure

Three caveats determine how far the conclusions in section 6 are entitled to go.

###### Synthetic targets

The benchmark recovers parameters of a model from data generated by the same model, so it measures the optimization problem in isolation, not the model–data mismatch one would face on real recordings; this is by design (section 4.1.4) but means the radii reported here are upper bounds on what the same setup achieves experimentally.

###### Single trace

A single 100 ms step current is not enough to pin down the model parameters; the problem is under-constraint. The multi-trace extension of section 4.1.3.3 is the natural address but lies outside the benchmark’s scope.

###### Fixed hyperparameters

The configurations in table S1 are the hyperparameters used in this single-stage setting; per-scenario, per-*δ* or per-method HP tuning could possibly shift radii up — this is part of the tuning problem which we will discuss in the next chapter (ch. 6). The radii reported here are therefore meant to characterize the methods as one would actually deploy them, not their best possible behaviour under unconstrained tuning.

#### 5.3.3 Subthreshold Recovery: Gradient Descent Without Spikes

The recovery-radius benchmark leaves an ambiguity: when a gradient method loses to Nelder–Mead, is that because gradient descent is the weaker optimizer, or because the surrogate gradient through the spike mechanism is a poor search direction? The subthreshold benchmark of section 4.1.5 settles this by removing the spike path entirely. Below rheobase only {*C*_*m*_, *E*_*L*_, *g*_*L*_} shape the trace, the custom-VJP surrogate never fires, and the MSE loss becomes a smooth three-parameter regression — the cleanest possible probe of gradient descent against a derivative-free baseline on identical, paired scenarios. The verdict reverses that of the spiking benchmark: with no spikes in the way, gradient descent is the stronger optimizer, not the weaker one.

Figure 11 shows the outcome. Gradient descent converges (clean-simulation MSE ≤ 0.1 mV^2^) on 80 % of the 200 scenarios against Nelder–Mead’s 53 %, and it holds this lead at every perturbation scale, from 100 % vs 70 % at *δ* = 0.1 to 56 % vs 46 % at *δ* = 0.4. The paired comparison is unambiguous: on the 89 scenarios where the two methods disagree on convergence, gradient descent is the one that succeeds in 71 of them and Nelder–Mead in only 18 (McNemar *p <* 10^−4^).

**Figure 11.**
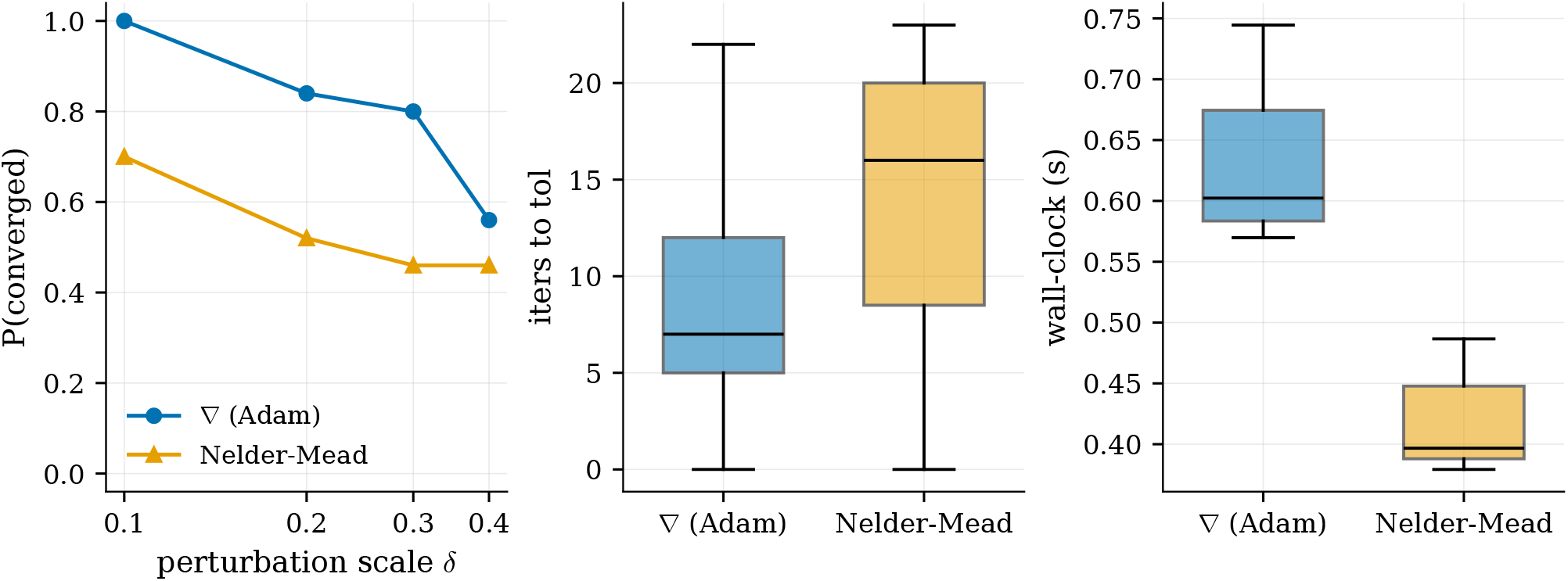
Subthreshold recovery, gradient descent (Adam) vs Nelder–Mead on the identical MSE loss, paired over 200 scenarios. (a) Probability of convergence (clean re-simulation MSE ≤ 0.1 mV^2^) as a function of the perturbation scale *δ*; gradient descent dominates at every *δ*. (b) Loss evaluations to reach tolerance and (c) wall-clock time per run, over the converged runs only. Gradient descent reaches tolerance in fewer forward-simulation evaluations — the cost axis that matters for fitting — even though a single evaluation carries a backward pass and so costs more absolute wall-clock time on this three-parameter problem.

Both optimizers are given the same budget of 25 model evaluations. On this axis gradient descent is decisively more sample-efficient: it reaches the convergence tolerance in a median of 9 fewer evaluations than Nelder–Mead (Wilcoxon *p <* 10^−4^, lower in 80 % of pairs), while also reaching a lower final loss (median final MSE 0.027 mV^2^ against 0.091 mV^2^; better in 77 % of pairs, *p <* 10^−4^). The gradient carries per-evaluation information that lets each forward simulation count for more, which is exactly the currency that matters once the forward model is expensive. Wall-clock time runs the other way on this three-parameter toy problem — the backward pass makes a gradient evaluation more costly than a bare loss evaluation, so at a matched evaluation budget gradient descent takes longer in absolute seconds — but that gap is an artefact of the trivial problem size, where autodiff overhead has nothing to amortize against, and it inverts as soon as the per-simulation cost dominates the per-step bookkeeping.

There is one important caveat, and it connects directly back to the loss geometry of section 5.2. Converging to a lower voltage MSE does *not* translate into better parameter recovery. Measured as bound-normalized parameter error, gradient descent is if anything slightly worse than Nelder–Mead (median 0.081 vs 0.052; better in only 24 % of pairs, and the paired test does not favour it), even though it wins decisively on the loss it actually minimizes (fig. S7). This is the subthreshold signature of the degeneracy already visible in the landscape plots: *C*_*m*_, *E*_*L*_ and *g*_*L*_ trade off along an extended low-loss valley (the *g*_*L*_–*E*_*L*_ direction of section 5.2.1), so many distinct parameter triples produce near-identical subthreshold traces. A better optimizer finds a lower point in that valley without necessarily landing closer to the true parameters. The subthreshold problem is therefore well-behaved as an *optimization* problem and simultaneously ill-posed as an *identification* problem — a distinction we return to in section 6.

Two conclusions carry forward. First, gradient descent with the Jaxley AdEx channel is not intrinsically broken: given a differentiable path to the loss, it is competitive with and often better than a strong derivative-free baseline, and more sample-efficient in forward-simulation evaluations. The failures in the spiking benchmark of section 5.3.2 are properties of the spike-mechanism gradient and the loss surfaces it has to cross, not of gradient descent as such. Second, even in this benign regime the loss and the parameters are only loosely coupled, which sets a ceiling on what any single-trace method — gradient or derivative-free can recover, and motivates the multi-trace direction of section 4.1.3.3.

## 6 Discussion

The last chapter gave an overview over the hard facts. But where does that leave us and what does this mean going forward? Section 6.1 sets out what the thesis contributes regardless of the optimization outcome. Section 6.2 then shows why gradient-based fitting failed and weighs the ways forward (section 6.2.4), including the alternative neuron models of section 3.2 that avoid the surrogate bias by construction. Lastly, we want to discuss the limitations of this thesis in section 6.3.

### 6.1 Simplified Neurons in Jaxley

Independently of the optimization results that occupy the rest of this chapter, the implementation contribution stands on its own. We extended `Jaxley`, still in early development, with a new AdEx channel^*^ (sections 4.3.1 and 5.1). More substantively, we were the first to try a surrogate-gradient mechanism for any of its simplified models for parameter fitting. The `custom_vjp` construction generalizes beyond AdEx: the same pattern — voltage integration inside `update_states`, surrogate Vector-Jacobian Product (VJP) on the spike indicator — applies verbatim to the pre-existing LIF and Izhikevich channels. There are still lots of movement and structural changes to be expected within the framework. The development team plans to publish a different surrogate gradient implementation with the `Jaxley` v1.0.0 release.

The practical leverage of this is broader than the gradient-fitting story `Jaxley` was originally built for. Differentiability of biophysical models is one of its headline features, but the same simulator — JIT-compiled, `vmap`-batched, GPU-capable — is equally useful for derivative-free pipelines that consume many cheap forward passes: evolutionary algorithms, parameter-space exploration as in Guarino et al. [19], and simulation-based inference [21]. The infrastructure is therefore valuable independently of whether the gradient consumer ultimately delivers on the recovery problem — an observation that becomes material in section 6.2, where a derivative-free baseline running on this same simulator outperforms the gradient methods it was originally built to support.

### 6.2 Differentiable AdEx for parameter optimization doesn’t work

The recovery-radius benchmark of section 5.3.2 gives an unambiguous and uncomfortable answer. Across all three scenarios and every perturbation scale, no gradient method matched the Nelder–Mead simplex on the same objective. MSE was actively harmful, finishing at or below the floor on every run. Guarino and Van Rossum both clear the floor at small *δ*, but their radius never exceeds *δ* = 0.1 — an order of magnitude short of Nelder–Mead. The answer to the thesis question follows directly: gradient-based optimization produces useful updates for the AdEx model only inside basins that are already small enough for derivative-free search to cover. This kills the argument that motivated the work. Gradient search is not a *press-one-button-fit-the-model-solution*. The order-of-magnitude speedups Deistler et al. [5] report for HH-type fitting were expected to carry over to simplified models once surrogate gradients were added; they do not. Gradient fitting works for HH models because their dynamics are smooth end to end. AdEx dynamics are not, and a surrogate patch at the threshold does not change that.

One objection is that the gradient methods lose because the soft Guarino features distort the objective in a way the simplex does not see. The benchmark rules this out. The two Nelder–Mead variants report identical recovery radii on every scenario. One runs on the soft Guarino features (`nm_guarino`), the other on exact ones (`nm_guarino_hard`), and they differ by at most one direction in thirty. The two families share the objective, so what separates them is the gradient consumer. The simplex treats the loss as a black box and reads its slope by polling the corners of its simplex. On a landscape of narrow valleys and wide plateaus it steps over the flat regions that stall gradient descent, and it never meets the pathologies that our stabilization toolbox (section 4.1.3) was built for. Gradient descent meets all of them. The toolbox only rescales gradient magnitude, but the pathologies that matter are about direction and existence, which rescaling cannot reach. Three mechanisms plausibly drive this, in decreasing order of evidential support.

#### 6.2.1 Failure mode 1: Surrogate gradients as biased estimators

The “approximate gradient” label is misleading. The true gradient *∂ ℒ/∂θ* runs through the Heaviside derivative and is zero almost everywhere, so there is nothing to approximate. The surrogate instead returns the exact gradient of a *different* objective: a smoothed loss 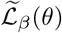 in which the hard threshold is replaced by a step of width ~1*/β* [10]. Its minima meet those of the true loss ℒ only as *β* → ∞.^*^ At any finite *β* the two are offset by a displacement of order 1*/β*. That displacement is the bias. Gradient descent reaches a minimum of 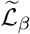 cleanly, but that minimum need not sit where the individual spike times require.

This bias stayed invisible in the setting where surrogate gradients were first validated. Spiking-network classification averages it away: the error is shared across many neurons and an underdetermined set of readout weights, so displacing any single weight barely moves the task loss [9, 16, 23]. Single-neuron AdEx fitting has no such averaging. The displacement lands directly on the fitted parameters 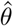 and shows up as a finite recovery radius, even though the gradient is non-zero and the loss keeps dropping.

#### 6.2.2 Failure mode 2: Loss-landscape pathologies differ per loss

The artefacts catalogued in section 5.2 share one property: they come from the loss formulation, not from the surrogate. Replacing the surrogate with an exact spike-time gradient would remove none of them.

For MSE, deleting a misaligned spike lowers the pointwise voltage error more than aligning it would, so the non-spiking trace is a genuine minimum of the loss (section 5.2.2). For Guarino, the missing-feature penalty puts a step in the forward loss at the boundary between no spikes and a few spikes. The validity gates smooth that step only locally, so the ridge that blocks descent (section 5.2.3) sits in the loss surface itself. For Van Rossum, a change in spike count is a property of the spike train *s*(*t*), and the kernel convolution acts downstream of it, so *K*\* *s* still jumps whenever a spike appears or disappears (section 5.2.4).

The three losses differ in how much softening they stack on top of the surrogate, and the recovery numbers of table 5 partly track this. Van Rossum adds the least: its kernel convolution and *L*^2^ norm are exactly differentiable, so the surrogate is its only source of bias. Guarino adds more, routing through soft validity gates with their own sharpness *β*_*v*_. If bias grew with softening layers, Van Rossum should beat Guarino everywhere — and it does: a tie on `tonic` (+1.00) and wins on `adaptation` (+1.00 vs +0.54) and `initial_bursting` (+1.00 vs +0.90). The least-softened loss carries the least bias, though both trail Nelder–Mead by an order of magnitude. MSE stays at the bottom throughout, which the non-spiking attractor above already explains.

Guarino also carries a tuning cost the others avoid. Four softness hyperparameters fix its landscape and its gradient direction: the feature weights, the missing-feature penalties, the validity-gate sharpness *β*_*v*_, and the surrogate *β*. All of them have to be set per cell. A derivative-free pipeline pays none of this, because exact thresholds and integer spike counts are valid feature definitions when no gradient is required.

The contrast with FM1 is clean. FM1 says the optimizer is pulled towards the minimum of the wrong objective. FM2 says the objective itself fights gradient descent, and no change to the surrogate repairs that.

#### 6.2.3 Failure mode 3: Vanishing gradients through long unrolled simulation

Suppose FM1 and FM2 were both fixed, giving a clean per-step surrogate and a friendly loss surface. The parameter gradient would still have to survive roughly 5000 multiplicative leak-Jacobian terms. Section 2.3.2 shows this is a severe vanishing regime for the AdEx parameters that shape subthreshold dynamics.

The usual fixes from the SNN literature do not fit here. Truncated BPTT adds a second approximation bias on top of the FM1 bias we are trying to avoid, because it cuts the backward pass after *k* steps and detaches it from older states. This stops the gradient vanishing, but it drops all credit assignment through dependencies longer than *k* steps — the truncated gradient is a biased estimate of the full one. That bias is separate from, and stacks on, the surrogate bias of FM1. Eligibility traces [29] need a per-step error signal that single-trace fitting does not provide. A larger step Δ*t* shortens the unroll but costs accuracy on the exponential spike term, exactly where the surrogate matters most.

This is the most speculative of the three modes. It acts at every gradient step together with FM1 and FM2, so the benchmark cannot isolate it.

#### 6.2.4 Synthesis and forward directions

The three modes act together at every gradient step. The surrogate bias sets up the displacement (FM1), the loss landscape decides whether the biased minimum can be reached (FM2), and the decay decides whether any signal survives the trip backward (FM3). The data cannot weigh them against each other, but the negative result holds whichever one dominates.

The cleanest way forward is not a better surrogate or a better loss, but a different neuron model that avoids the bias by construction. The Klos–Memmesheimer construction [24] from section 3.2 does this. It gives a closed-form between-spike solution for the QIF family and a pseudospike mechanism that keeps silent neurons trainable. Our methods descend on a smoothed loss and hope it tracks the true one; theirs descend on the true loss directly.

Which path is right depends on what the pipeline is for. If exact spike times matter more than fidelity to a particular biophysical model, that construction is a better starting point than more surrogate tuning on AdEx. If the AdEx biophysics are required, the bottleneck is the loss landscape and the gradient magnitude, not the optimizer.

On the landscape side, the untried options are multi-trace constraints (section 4.1.3.3), curriculum schedules, hybrid objectives that pair Van Rossum’s kernel smoothness with feature-based timing terms, and signal-processing spike-train distances.

On the magnitude side, the stabilization toolbox of section 4.1.3 only rescales the signal that already survives, so the useful interventions attack the source instead. Forward-mode differentiation carries the sensitivity *∂V/∂θ* forward alongside the state in a single sweep, cheap here because AdEx has only a handful of parameters.^*^ It does not change the total gradient, since forward and reverse mode return the same derivative, and unlike truncated BPTT or eligibility traces it adds no approximation of its own. What it does add is a sensitivity at every timestep, so a loss that reads out error along the whole trace can supply credit locally instead of pushing one signal back through the full unroll (section 6.2.3). Per-parameter preconditioning targets the magnitude asymmetry of section 5.2. Because *b* and *τ*_*w*_ act only through spike events or slow relaxation, their gradients arrive far below those of the continuously-acting parameters, and a shared step size either starves them or overshoots the rest. Rescaling each update by its own gradient magnitude puts the axes on a common footing, and Adam’s second-moment term already does a weak version of this. Neither fix is unconditional. Forward-mode only relieves the FM3 decay when the loss is distributed in time, which is the same per-step error signal the eligibility traces of section 6.2.3 require. A voltage MSE supplies it, but that loss fails for the FM2 reasons above, and the spike-timing losses we would rather use do not supply it at all. Preconditioning in turn helps only where the small magnitude is a scaling artefact, not a gradient that is structurally absent, as happens for *b* on a silent trace.

### 6.3 Limitations

The negative result above is the outcome of a benchmark whose scope is deliberately narrow. Five constraints in particular bound the conclusion’s magnitude.

#### Single model

The recovery-radius benchmark is on AdEx only. The bias argument of section 6.2.1 predicts that gradient methods should fare worse on LIF (slope zero at threshold, surrogate bias maximal) and somewhere between AdEx and QIF on Izhikevich; neither prediction is tested. Substituting `FireSurrogate` (or an Izhikevich equivalent) into the benchmark pipeline is a drop-in change and is the cleanest way to falsify the surrogate-bias view.

#### Single stimulus protocol

All scenarios use 100 ms step currents. Time-varying or ramped inputs might exercise gradient pathways through *τ*_*w*_ that step currents cannot, and the multi-trace setup of section 4.1.3.3 is implemented but not used in the headline benchmark. A multi-trace replication of the recovery-radius experiment is the natural extension and would speak directly to the adaptation-parameter identifiability question deferred in section 4.1.4.2. However, parameter recovery on a single trace for small perturbations will almost certainly be a necessary precondition for any successful multi-trace optimizations.

#### Fixed hyperparameters per method

Each method runs with one configuration across all scenarios and perturbations. No per-cell tuning, no architecture search. Guarino’s hyperparameter brittleness (section 6.2.2) is itself evidence that an Hyperparameter (HP)-search budget could shift the gradient-method radii up; we have not quantified by how much, and this is the most likely angle from which a follow-up could partially overturn the result. However, if scenario specific hyperparameter tuning is needed, to successfully run this method, one could directly sample over the initial parameter space using the same computational resources.

#### Failure modes as candidates, not isolated causes

The three failure modes are entangled at every gradient step. Isolating FM1 cleanly requires the LIF substitution above; isolating FM3 requires sweeping simulation length while holding FM1 and FM2 fixed; isolating FM2 requires an exact-gradient method on AdEx, which is the construction that does not exist for this model family. The chapter’s negative conclusion is robust to whichever component dominates, but the diagnosis itself is not substantiated by isolation experiments.

#### Synthetic targets

The benchmark uses model-generated data, so it measures the optimization problem in isolation rather than the model–data mismatch one would face on real recordings. The recovery radii reported are upper bounds on what the same setup would achieve experimentally. Since no perfect recovery can be expected when fitting to real neurons, a (further) drastic drop in fit quality can be expected.

## 7 Conclusions and Future work

Simplified neuron models remain a key tool for large-scale biophysical brain simulations. To fit those models efficiently we aimed to replace sampling-based optimization using a surrogate gradient construction to enable gradient-based algorithms. We successfully implemented the AdEx model using the Jaxley framework and contributed it to the project. We matched runtime performance of highly optimized, target-compiled C++ code of a Brian2 reference implementation and performed ~ 200× faster than the default Python implementation. Because `Jaxley` leverages `JAX`, the model supports CPU, GPU and TPU as target platforms without any code changes. This powers large-scale simulations and model-fitting applications regardless of the choice of optimizer.

Gradient-based fitting did not match the derivative-free counterpart across 2700 runs in any of the spiking regimes. Our systematic benchmark showed that gradient-based fitting is currently not viable for AdEx, even with the state-of-the-art stabilization techniques used for HH-model fitting. This verdict comes with one revealing exception: within the subthreshold regime, gradient-based fitting outperformed derivative-free Nelder-Mead with a convergence rate of 80% vs 53%. This indicates that propagating gradient information through the spike mechanism remains the key challenge.

We investigated the resulting challenges when adapting loss functions from derivative-free methods to gradient-based methods. One important finding is that biases introduced by softness parameters do not seem to affect derivative-free methods. However, the shape of the error surface can affect gradient-based methods more due to ridges or missing gradient information in regions where the neuron is silent. Specifically, voltage-based losses like MSE fail structurally. MSE prefers flat, silent traces to slightly misaligned spikes, which makes it unusable for spike-train fitting. Feature-based losses escape that problem but introduce design challenges and complications like error surfaces with hard ridges that block gradient descent or many settings that require hand-tuning.

Section 6.2.4 already mentioned investigating the use of different (differentiable) neuron models. In the context of surrogate-gradient based optimization, this would allow us to better understand the contribution of surrogate biases.

A second very interesting future work topic would be designing and evaluating different loss functions. Investigating efforts that might have been established in signal processing, different areas of Machine Learning (ML), or other research areas could lead to interesting transfer applications. Better understanding the effect of loss functions when gradient-based optimization works well (e.g. in the context of using HH-type models) might provide valuable insights. In the context of feature-based losses we mentioned the problem of too many hyperparameters. This is made worse when taking feature weighting into account — this is a problem of multi-objective optimization. Pareto fronts have not been explored or discussed in the context of this work due to time constraints.

Analysing different stabilization techniques to reduce vanishing/exploding gradients could most likely improve the qualitative results of this method. Using multi-trace optimization could help with the under-constrained fundamental problem of fitting the AdEx model to spike timings, which is an essential next step in the task of neuron model fitting.

## ACKNOWLEDGEMENTS

We thank Erik Fransén for both serving as the examiner of the Master’s thesis on which this work is based, and for his valuable comments and feedback on the manuscript itself.

## AUTHOR CONTRIBUTIONS

P.M. designed and implemented the method, performed the experiments and wrote the manuscript. A.K. proposed the topic, supervised the project and revised the manuscript.

## COMPETING FINANCIAL INTERESTS

The authors declare no competing financial interests.

## A Figures and Tables

### A.1 Detailed Experiment Configuration

**Table S1**

**Table S1.**
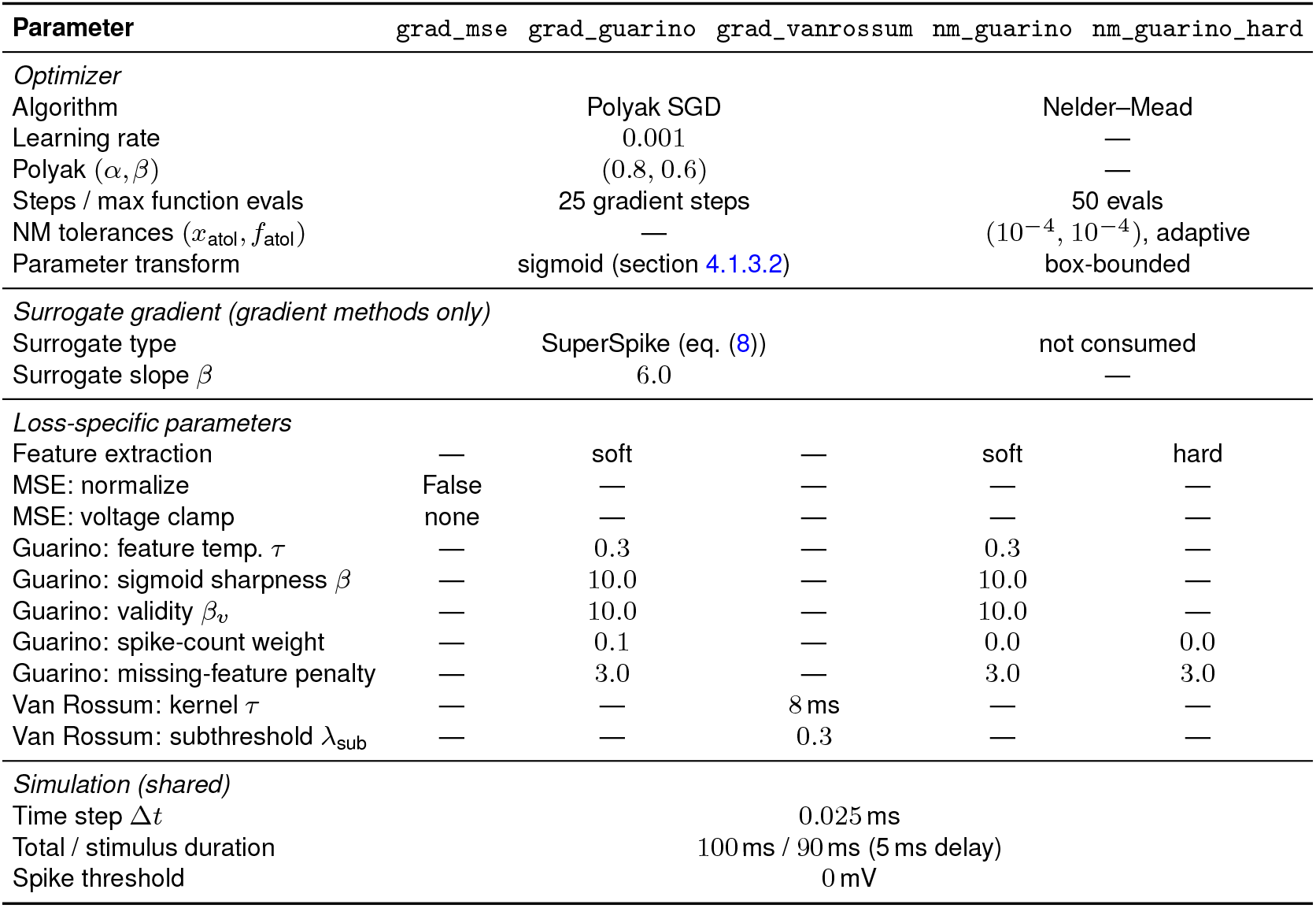
Method configurations for the recovery-radius benchmark. The three gradient methods share an identical optimizer, surrogate, and parameter-transform setup; they differ only in the loss function. `nm_guarino` optimizes the same soft Guarino loss with Nelder–Mead, evaluating but not consuming the surrogate gradient. `nm_guarino_hard` optimizes the same loss composition with *hard* feature extraction (threshold-crossing spike detection on both simulated and target traces), so the soft-feature hyperparameters *τ*, *β, βv* vanish from its configuration.

### A.2 Loss Landscape Analysis

To characterize the optimization problem independently of any particular training run, we computed each of the four loss functions on a dense two-parameter grid (20 × 20 = 400 points per pair) for 28 parameter pairs, for a total of 11,200 simulations and ~3.8 h of compute. The remaining parameters were held at the default initial values of table 2.

### A.3 Additional Results Figures

Supporting figures for section 5.

**Figure S1.**
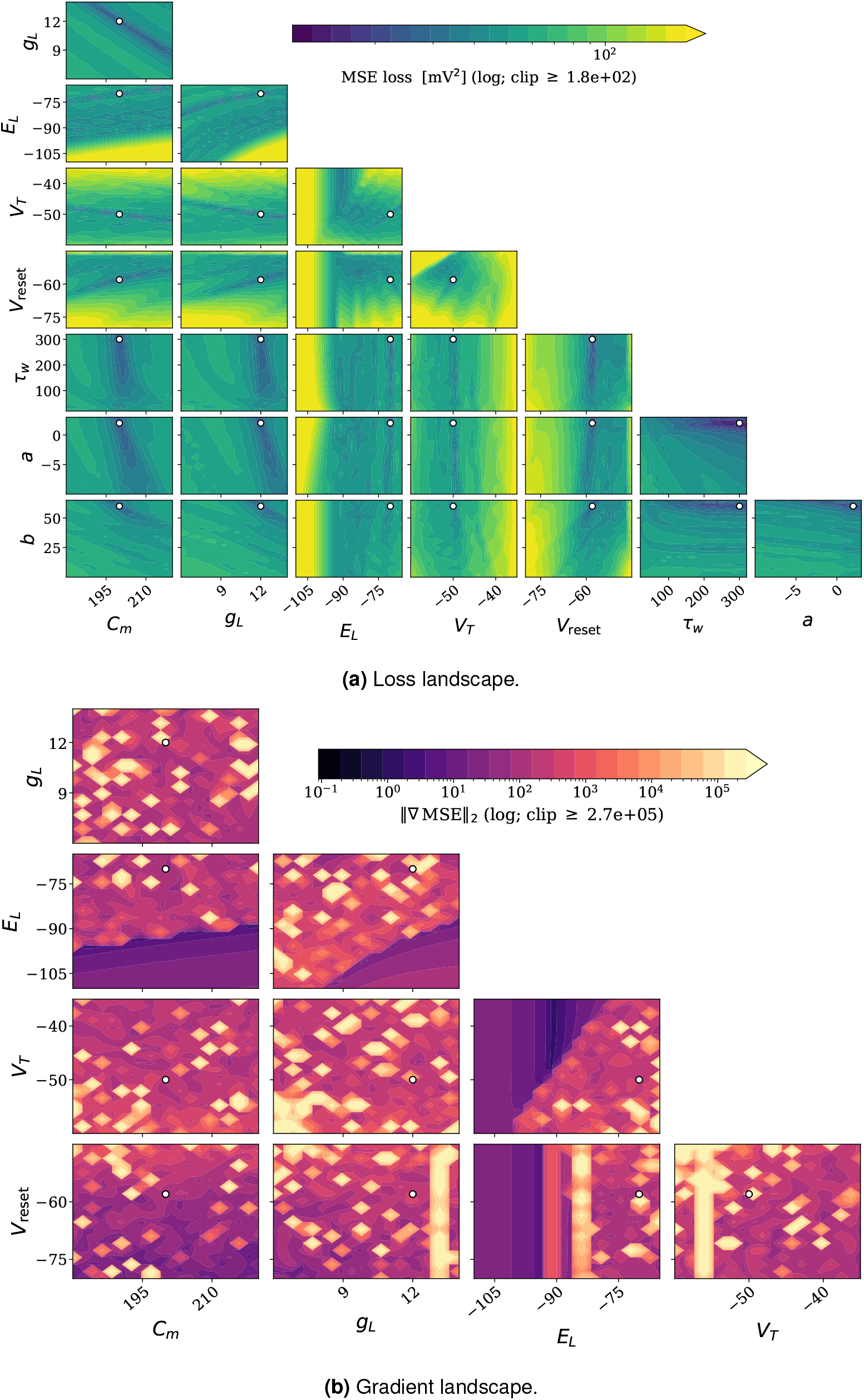
The MSE loss landscape — even in these simplified conditions where only two parameters at a time are changed — is characterized by lots of local minima. Some parameter combinations are also characterized by long shallow valleys.

**Figure S2.**
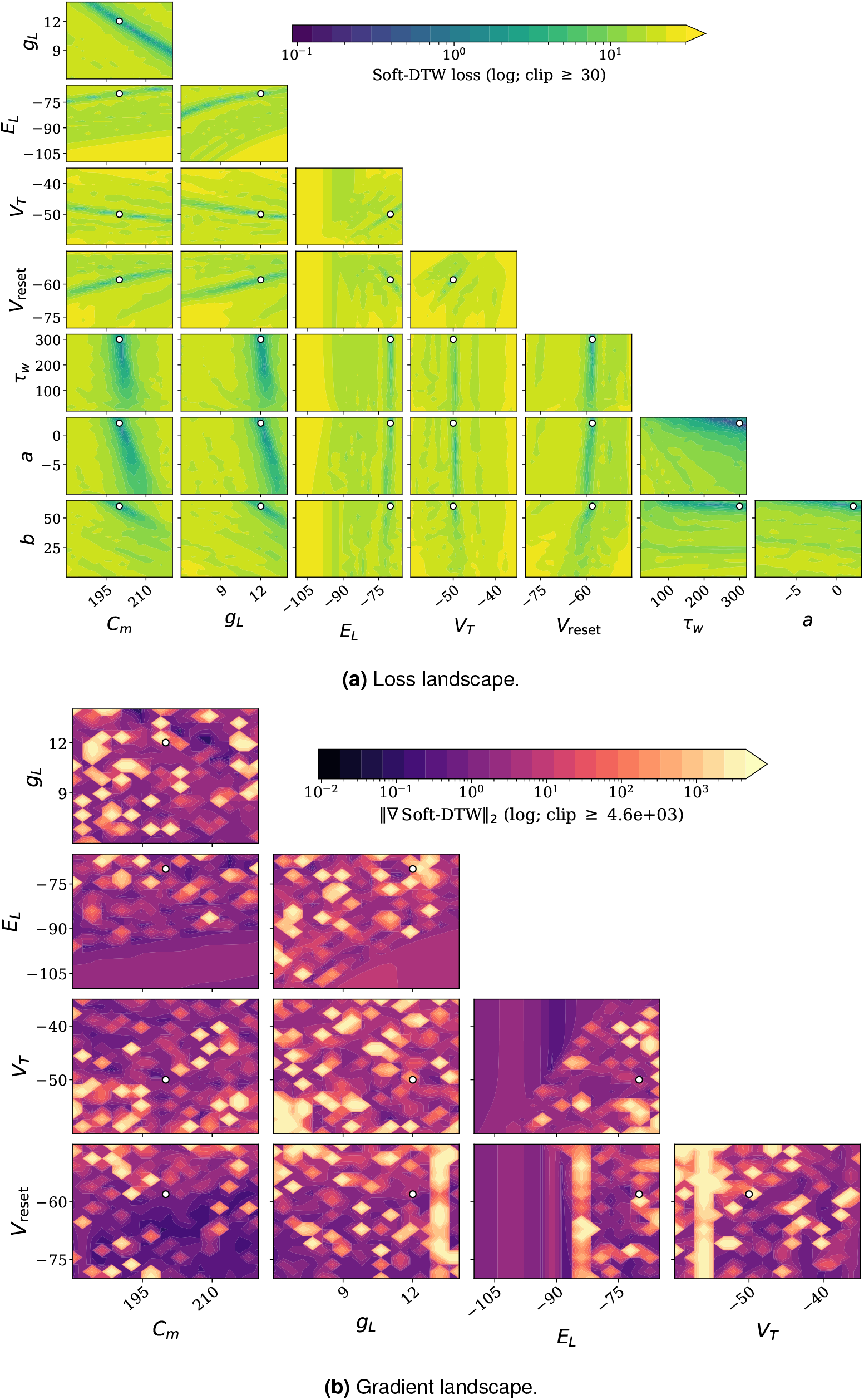
When observing the SDTW loss landscape, we identify very thin and deep valleys containing the global minimum. This correlates with the observation, that SDTW was only able to recover very small parameter perturbations. For larger perturbations, gradients were close to 0, which can be explained by the large loss plateaus visible.

**Figure S3.**
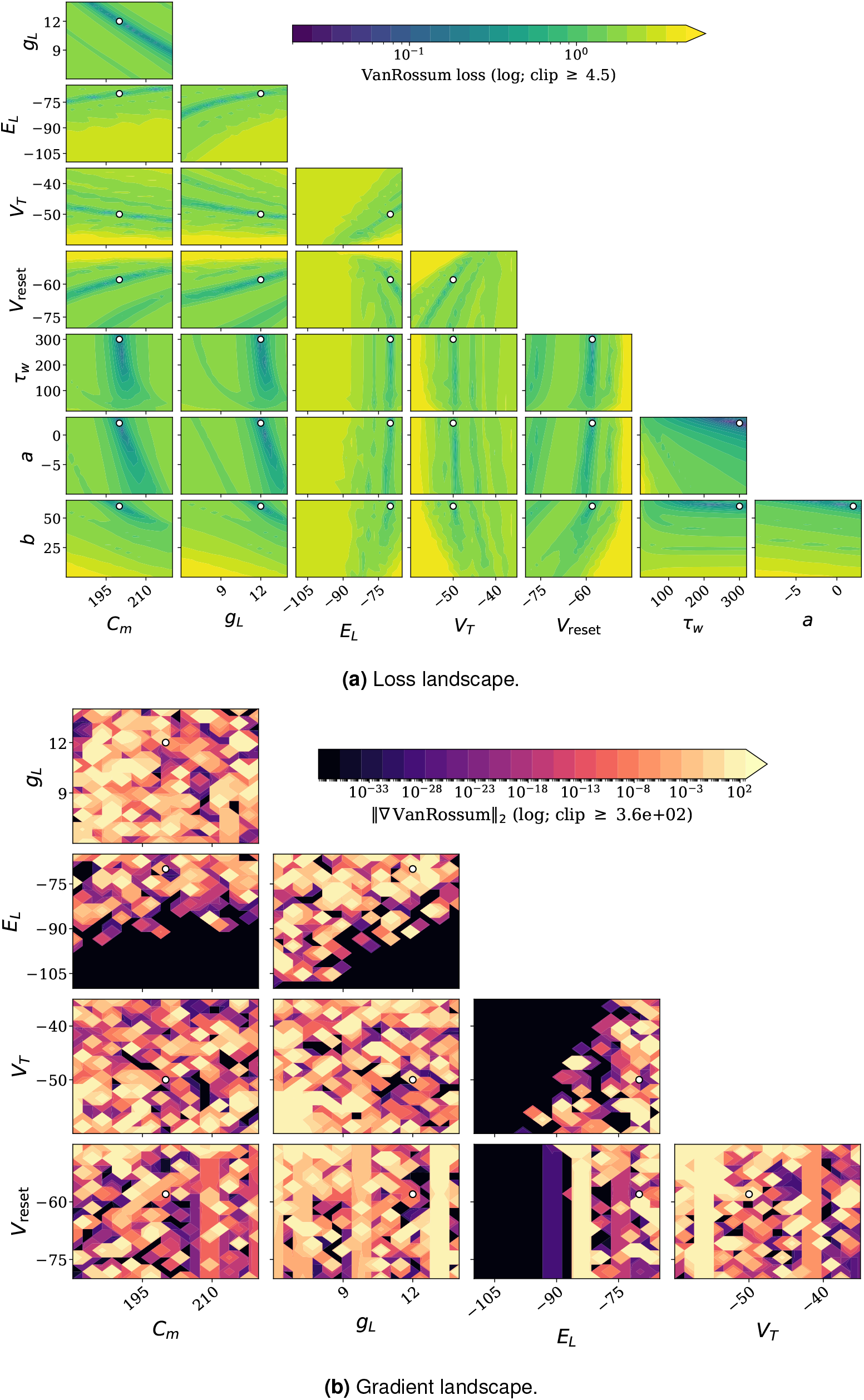
The Van Rossum loss promises to be a more interesting candidate based on these loss landscape plots. For example, the local minima are less pronounced than in the MSE landscape, even though still present. The global-minimum valleys tend to be less narrow compared to SDTW; however, they still tend to be surrounded by large loss plateaus.

**Figure S4.**
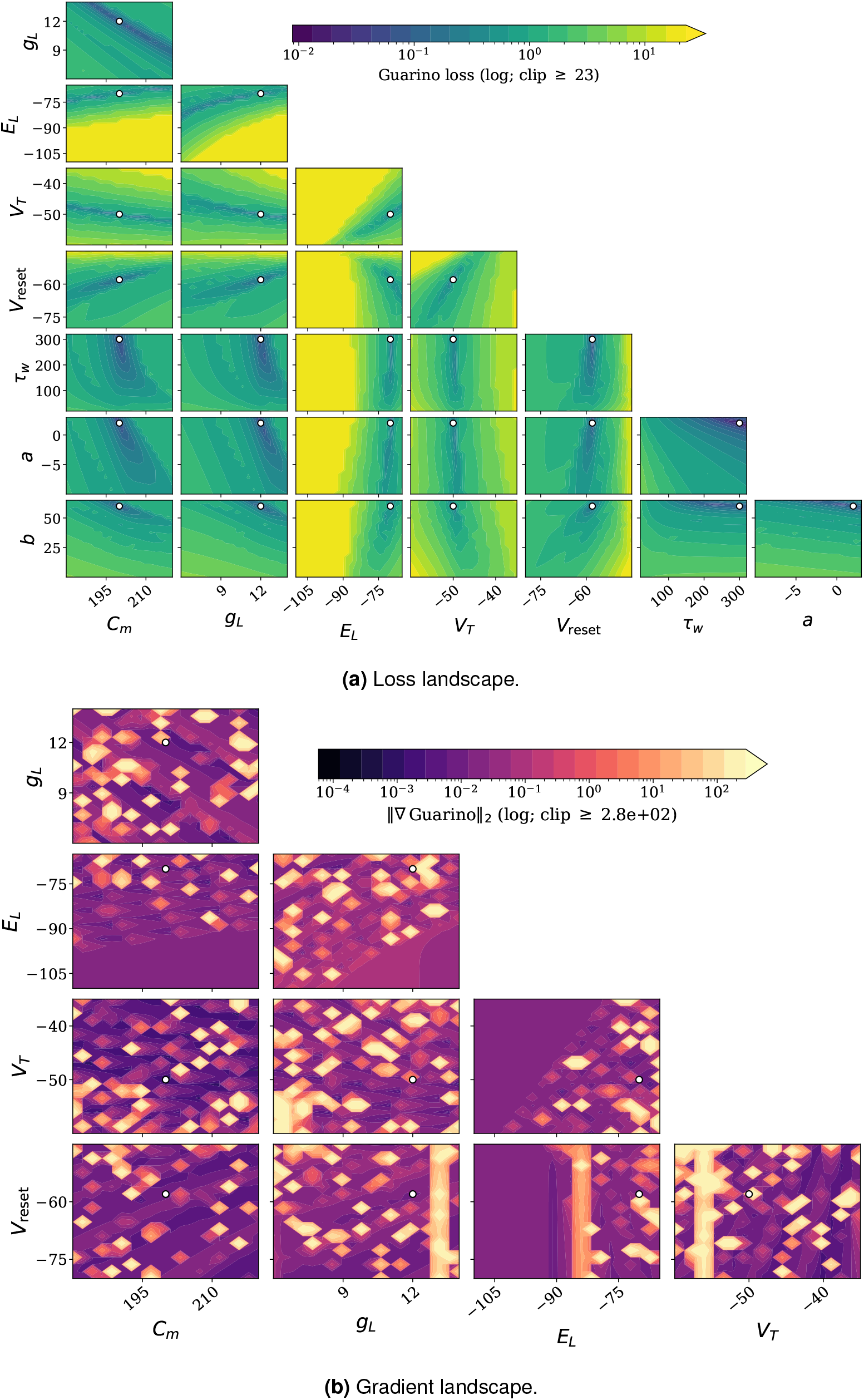
The feature-based loss shares lots of similarities with the Van-Rossum loss. It seems to be the most resistant against local minima in this testing configuration. However, it shares the large loss plateaus which could indicate small-to-zero gradients for ill conditioned initial parameters.

**Figure S5.**
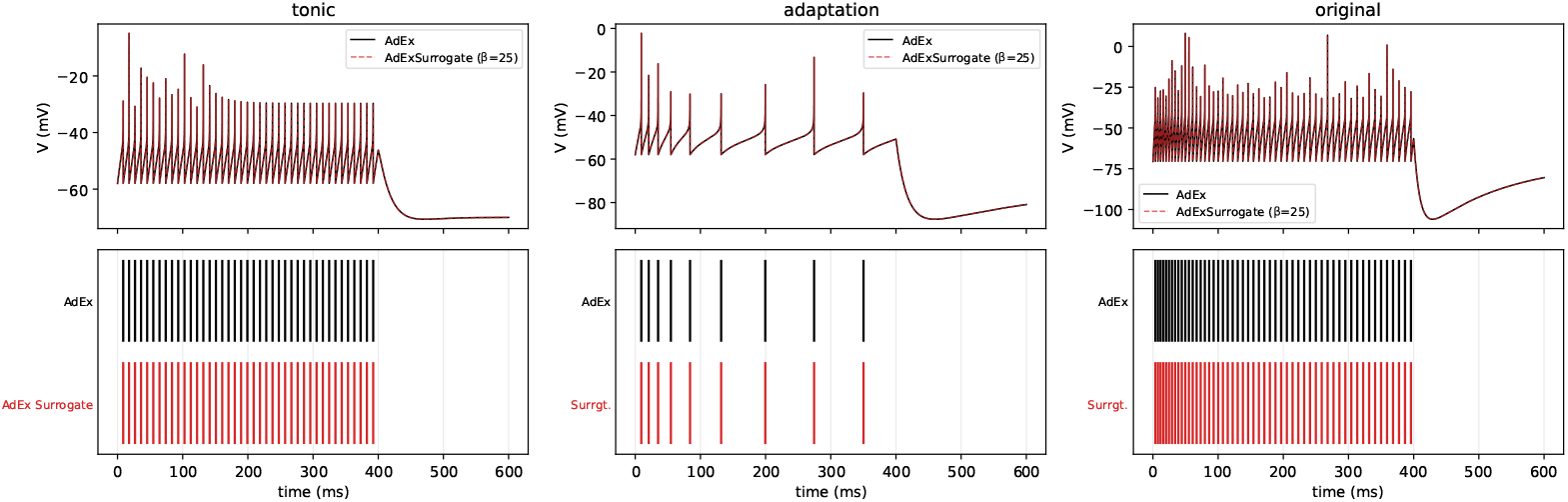
Comparison of `AdEx` (black, solid) and `AdExSurrogate` (red, dashed; sigmoid surrogate, *β* = 25) on the three Naud parameter sets. Top row shows membrane potential over time. Bottom row compares the corresponding spike trains. Both channels produce identical voltage traces and identical spike times as expected.

**Figure S6.**
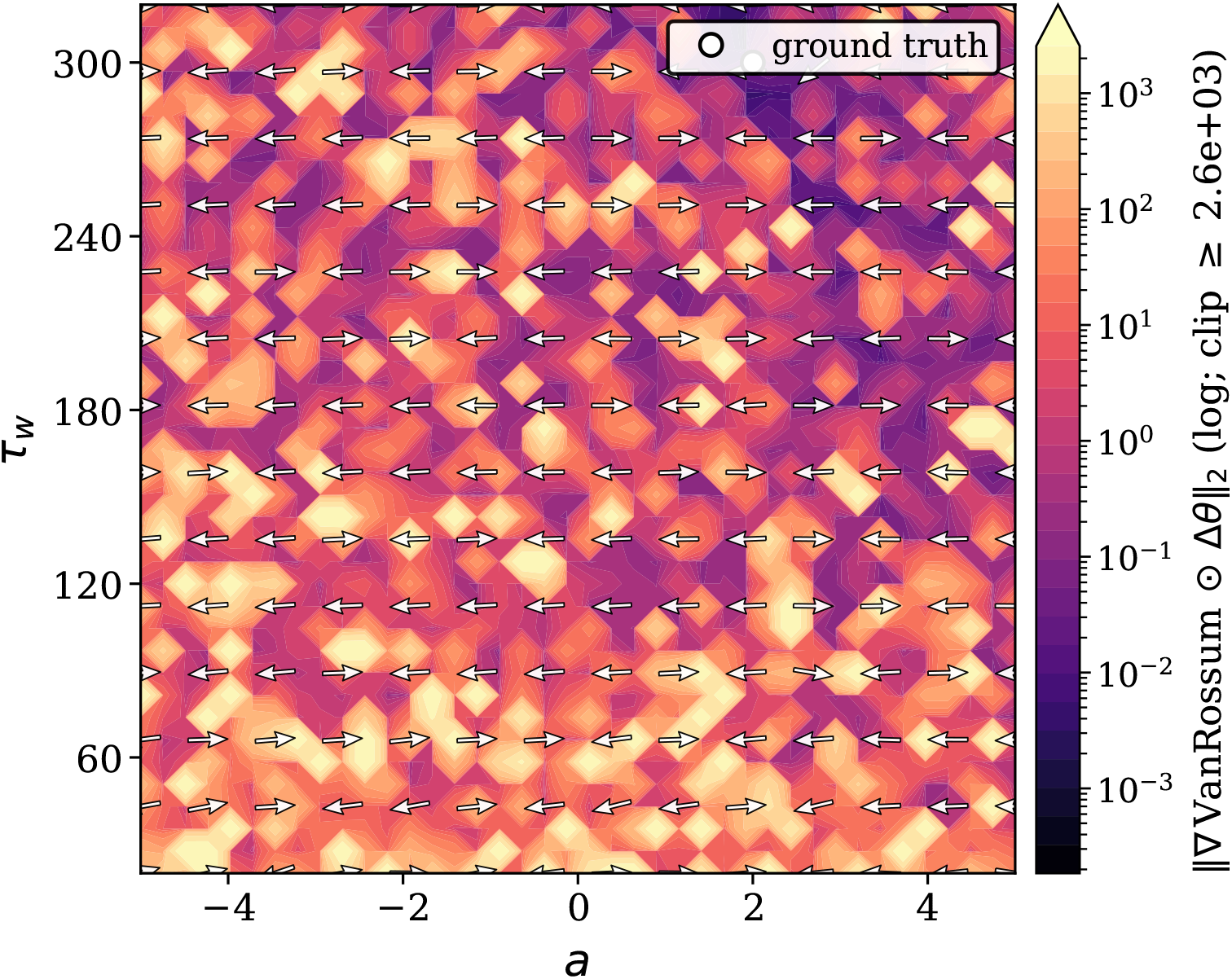
Van Rossum bound-normalized gradient field on the *a*–*τ*_*w*_ plane in the `adaptation` scenario (ground truth *a* = 2 nS, *τ*_*w*_ = 300 ms). SuperSpike surrogate, *β* = 10. The colour scale spans six orders of magnitude in bound-normalized sensitivity, but the quiver is dominated by the horizontal component throughout: |*∂*ℒ*/∂a* · Δ*a*| *≫* |*∂*ℒ*/∂τ*_*w*_ · Δ*τ*_*w*_ | across the whole slice.

**Figure S7.**
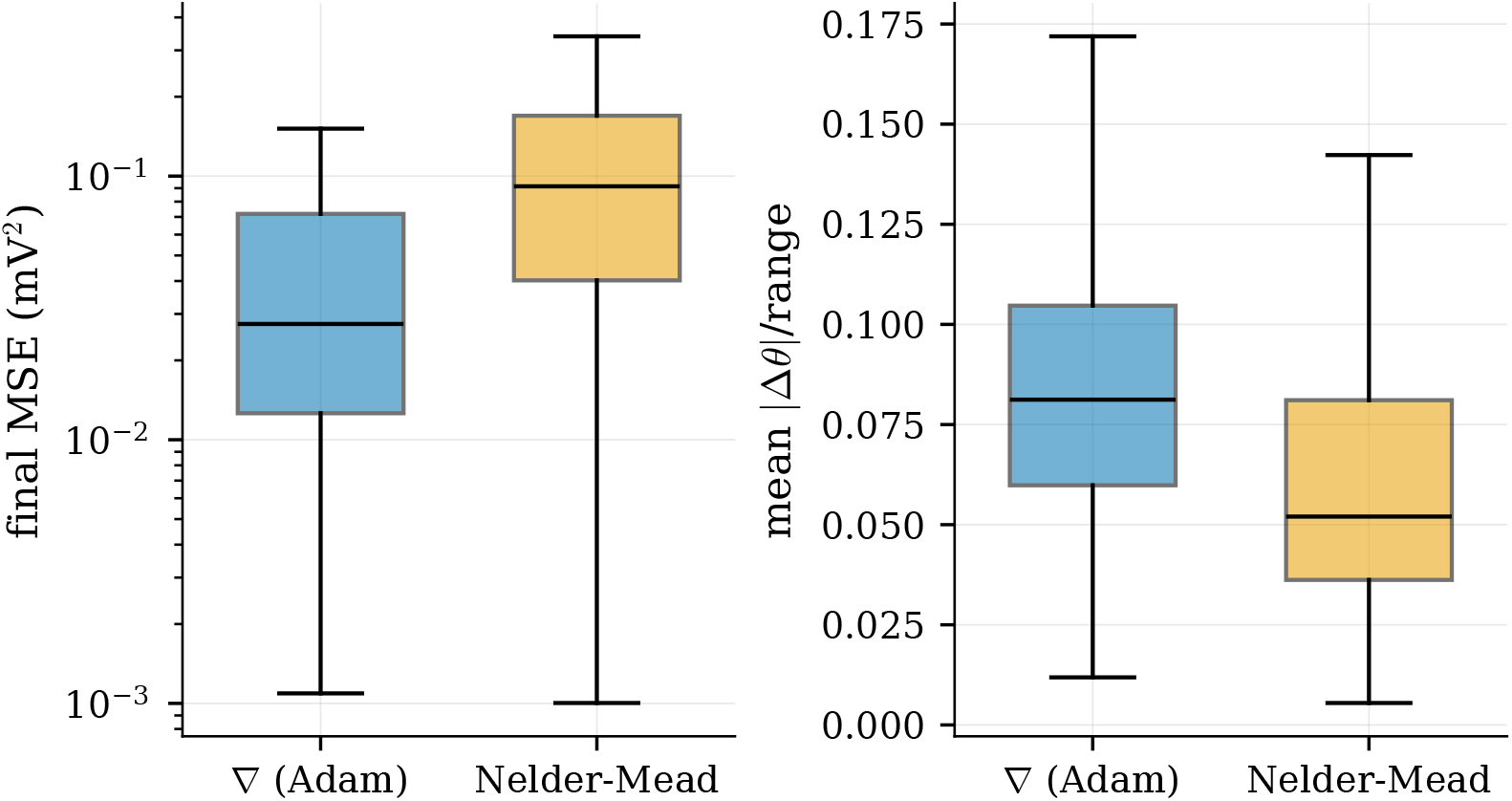
Fit quality over all subthreshold runs. Left: final MSE on a clean re-simulation (log scale) — gradient descent reaches lower loss. Right: mean bound-normalized parameter-recovery error — the two methods are indistinguishable, with gradient descent if anything slightly worse. Lower loss does not imply better parameter recovery: the subthreshold loss is degenerate in {*C*_*m*_, *E*_*L*_, *g*_*L*_}.

## Footnotes

* This is the equivalent to the inverse temperature *β* from the softmax function (eq. 10)

* “Mass” here is borrowed from measure theory / signal processing. A spike train is treated as a sum of point events on the time axis, and each event carries a mass—the integer or real number it contributes to integrals/sums of the trace.

† The adaptation variables are technically trainable exactly like all other parameters. However, as we will discuss in section 6, the gradient signal was too small and noisy for us to achieve sensible results for these parameters.

* Brian2 uses the same numerical integration scheme for both ODEs, whereas our AdEx implementation uses forward Euler for the voltage equation and exponential Euler for the adaptation equation. We mitigate an effect of this difference by choosing a small enough time step.

* We were able to contribute it to Jaxley, see PR

* The minimum shifts even though the kernel is centred on the threshold in voltage. Two effects combine. The smoothing acts inside the dynamics, in voltage, whereas the quantity being minimised is the parameter vector, so a smoothing centred in voltage is not centred in parameter space. And convolving an asymmetric loss valley with a symmetric kernel moves its minimum toward the gentler wall regardless, by an amount that grows with the width 1*/β*.

* Forward-mode cost scales with the number of parameters, so this advantage is specific to the low-dimensional single-neuron setting. On a large multi-compartment network reverse-mode BPTT is cheaper, and the binding constraint becomes trajectory-storage memory rather than the FM3 decay.

## Notes

https://github.com/paulmyr/Surrogate-Gradients-for-Gradient-Based-Parameter-Estimation-in-Simplified-Neuron-Models

### Competing Interest Statement

The authors have declared no competing interest.

